# Role of *N*-glycosylation as a determinant of ATG9A conformations and activity

**DOI:** 10.1101/2025.01.17.633517

**Authors:** Matteo Lambrughi, Mattia Utichi, Henri-Baptiste Marjault, Christian B. Borg, Sergio Esteban Echeverría, Kenji Maeda, Nicholas M.I. Taylor, Elisa Fadda, Marja Jäättelä, Elena Papaleo

## Abstract

In this study we investigate the effects of glycosylation at position N99 on the structural dynamics and lipid scrambling activity of ATG9A, a key autophagy protein, using microsecond all-atom molecular dynamics (MD) simulations. ATG9A is an integral membrane protein involved in autophagosome biogenesis, and glycosylation at N99 is known to play a critical, yet poorly understood role in its function. The MD simulations revealed that the hydrophilic central cavity of ATG9A supports lipid reorientation and partial transbilayer movements, consistent with its lipid scrambling activity observed experimentally. N-glycosylation at N99 was found to enhance cooperative interactions between protomers, facilitating lipid insertion and traversal within the central cavity. These findings align with the proposed mechanism of ATG9A role in lipid redistribution across the phagophore membrane during autophagy. However, mutagenesis experiments that abolish N-glycosylation in ATG9A (ATG9A^N99A^ and ATG9A^N99D^ mutants) did not show a significant change in autophagy flux, suggesting that further experimental approaches, such as lipid scramblase assays, are needed to pinpoint the function of glycosylation. In this study we also observed an asymmetric protomer conformations in ATG9A, contrasting with symmetric structures obtained from cryo-EM, suggesting that the structural heterogeneity of the protein could be further explored in cryo-EM datasets. Overall, the study highlights the importance of incorporating glycosylation in computational studies of membrane proteins and offers valuable insights into the molecular mechanisms of lipid transport in autophagy, with potential implications for other lipid scramblases and flippases.

## Introduction

Autophagy is a complex, evolutionarily conserved process through which cells degrade and recycle their components. This mechanism is essential for maintaining cellular homeostasis and responding to various cellular stresses^1^. Among the many post-translational modifications that modulate protein function, N-linked glycosylation (or N-glycosylation) plays a pivotal role in enhancing structural stability, protein folding, trafficking, and quality control ^2^. Additionally, it has been shown to regulate autophagy by modulating the function of autophagy-related proteins. Indeed, N-glycans are required for the chaperone-mediated autophagy function of the lysosomal membrane protein LAMP-2A ^3^.

Similarly, ATG9A (UniProt Q7Z3C6), another N-glycosylated autophagic protein, is essential for the formation of the double membrane vesicle known as the autophagosome. The autophagosome engulfs and delivers cargo to the lysosome for degradation. ATG9A is the only integral transmembrane protein in the autophagy machinery. During resting conditions, it traffics through different cellular compartments, such as the trans-Golgi network and endosomes ^4^. Upon autophagy induction, ATG9A is transported via vesicles to the phagophore assembly sites, where it acts as a seed for phagophore biogenesis and elongation ^5–10^.

ATG9A interacts with ATG2 proteins, which are lipid transfer proteins^11–13^ that form a heteromeric complex critical for phagophore biogenesis ^14,15^. The current evidence suggests that ATG2 proteins transfer lipids from the ER to the nascent phagophore. ATG9A, which has a lipid scramblase activity, redistributes lipids between the two leaflets of the phagophore membrane, enabling its elongation and the biogenesis of the autophagosome^13,16–21^. Human ATG9A and yeast Atg9 share a homotrimeric structure with a unique fold and a complex network of branched cavities (**Figure 1A-B**) ^18–20^. This network of cavities includes: i) a central cavity whose interfaces are provided by each of the three protomers, ii) a lateral cavity for each protomer, connecting the central cavity to the interface between the cytosolic leaflet of the membrane and the cytosol, and iii) a perpendicular cavity for each protomer, connecting the lateral cavity to the cytosolic side of the protein (**Figure 1A-B**). These cavities can provide different routes for lipid transfer between the two membrane leaflets and may play a role in lipid scrambling.

**Figure 1.**
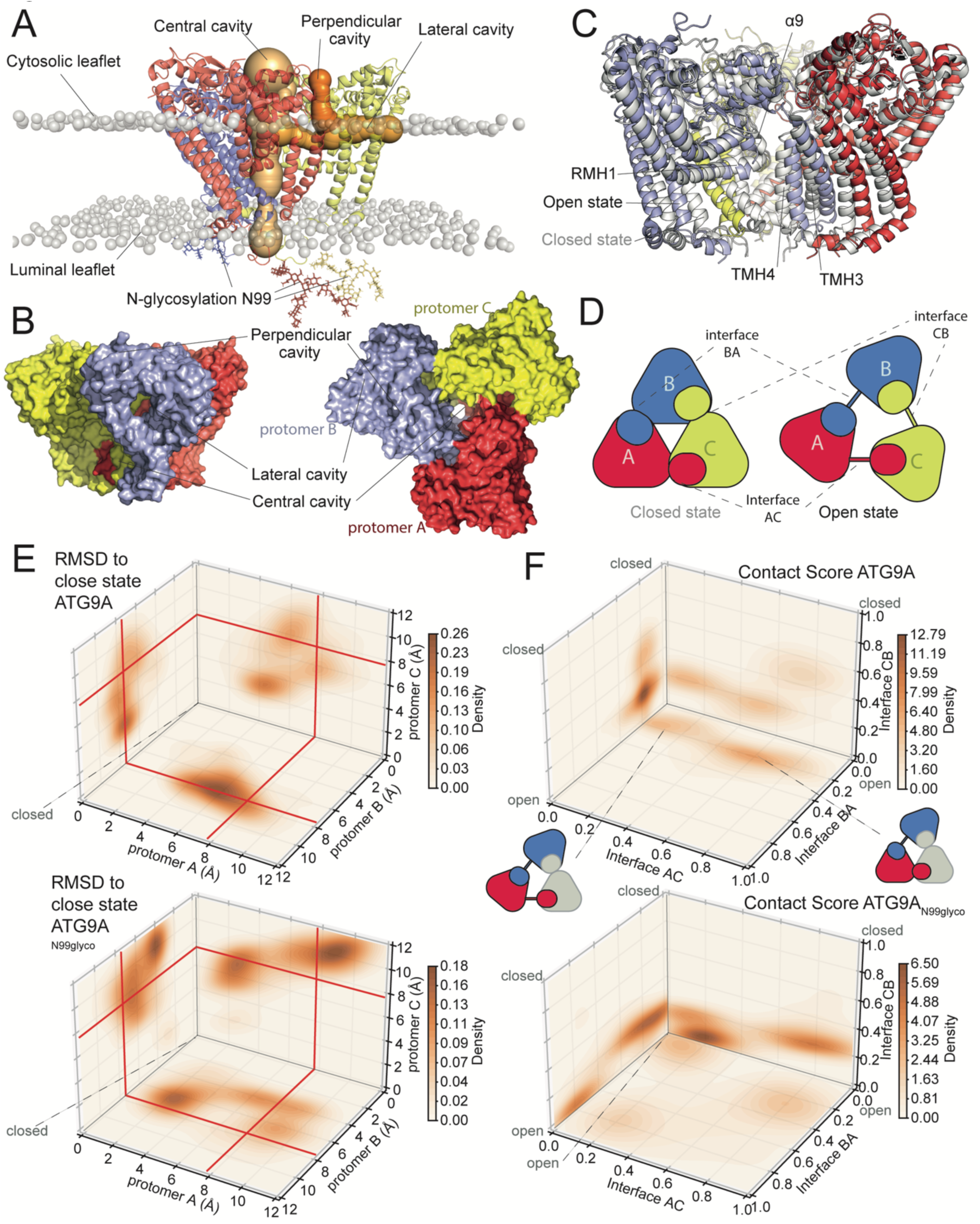
ATG9A homotrimer undergoes asymmetric closed-open conformational changes in the non-glycosylated and glycosylated state. (A) Starting model for MD simulations of *N-*glycosylated ATG9A homotrimer (ATG9A_N99glyco_). ATG9A is depicted as a cartoon representation, highlighting its three distinct protomers A (red), B (blue) and C (yellow). ATG9A has a complex network of branched cavities, including a shared central cavity and individual lateral and perpendicular cavities for each protomer. For protomer C, the radius of the internal cavities is visualized as orange spheres. The N99 *N*-glycosylation are represented as sticks. The lipid headgroups of the lipid bilayer are shown as gray spheres. (B) Surface representation of ATG9A homotrimer in the closed state shown from the side and top in the left panel and right panel, respectively. The network of cavities is shown and the protomers A, B, and C are highlighted in red, blue, and yellow. (C) Cryo-EM structures of ATG9A as cartoon representation, showing two distinct symmetric conformations of ATG9A protomers denoted as open (PDB ID 6WQZ) and closed (PDB ID 7JLP) states. In the open state, the protomers A, B, and C are colored red, blue, and yellow, whereas in the closed state, they are represented in white. These conformations are characterized by the different distance between the domain-swapped transmembrane helices (TMH3-4) and the reentrant membrane helix 1 (RMH1)-turn-helix α9, as well as the rest of the protomer and the TMH3–4 helices of adjacent protomers. (D) Schematic representation showing the open and closed symmetric conformations of ATG9A at the domain-swapped interfaces between the protomers (specifically at the AC, BA, and CB interfaces, named after the involved protomers). (E) Root Mean Square Deviation (RMSD) of the Cα atoms in the domain-swapped transmembrane helices of each protomer of ATG9A (upper panel) and ATG9A_N99glyco_ (lower panel) in relation to the experimental cryo-EM closed state for ATG9A. The plots show the bi-dimensional density projections of RMSD values calculated for the concatenated trajectories. For context, the RMSD between the closed and open state is around 7.8 Å, highlighted by red lines in the plots. Starting from the closed-state conformation, both ATG9A and ATG9A_N99glyco_ exhibit conformational changes towards the open state in at least one protomer. (F) Contact score values calculated for ATG9A (upper panel) and ATG9A_N99glyco_ (lower panel). The contact score quantifies the occurrence of protein-protein contacts between the domain-swapped TMH3-4 domains of each protomer and other regions of ATG9A that are present only in the cryo-EM structure of the closed state. A score of 1 indicates a close resemblance to the closed state. The plots show the bi-dimensional density projections of the contact score values of each protomer interface (AC, BA, CB) calculated for the concatenated trajectories. Both ATG9A and ATG9A_N99glyco_ assume asymmetric open states of the protomers, with the closed-to-open conformational changes driven by the separation of the domain-swapped interfaces of the protomers.

*In vitro* and *in vivo* experiments have demonstrated that human ATG9A is N-glycosylated exclusively at residue N99, making it the only occupied site among the four potential *N*-glycosylation sites (N99, N129, N224, and N507) in ATG9A^4,22^. Earlier studies indicate that the glycan at N99 is complex^4^, which is compatible with the localization of the protein in the trans Golgi.^9^

It has been suggested that the N-glycosylation of ATG9A contributes to its subcellular trafficking ^4,22^. The deletion of the region spanning residues 593–761 in the C-terminus of ATG9A results in a failure to traffic to the Golgi. Specifically, the LxM-type motif (residues 711– 713) has been implicated in the transport of ATG9A from the endoplasmic reticulum to the Golgi apparatus, as mutations in this motif reduce the amount of ATG9A forms with complex-type glycans. Additionally, an ATG9A mutant lacking residues 233–252 exhibits reduced ATG2A binding and autophagy flux while retaining scramblase activity ^15^. Despite the absence of complex-type *N*-glycosylation, this mutant still localizes to the Golgi when stably expressed in cells. This could be explained by the essential role of the C-terminal regions of ATG9A in both Golgi trafficking and maturation of the complex glycan at the N99 site. Although glycan maturation is impaired in the 233–252 deletion mutant, its Golgi localization remains unaffected, suggesting that ATG9A trafficking might follow a different modus operandi. One hypothesis is that ATG9A may act as a reservoir, ready to be rapidly mobilized to its final localization in response to specific stimuli ^4,22^.

N99 is located in a short loop between the first and the second transmembrane helices (TMH1 and TMH2) of ATG9A. It is positioned near the luminal opening of the central cavity and faces the domain-swapped TMH3 and 4 helices (**Figure 1A-C**). This loop remains unresolved in the electron density from all currently available cryo-electron microscopy (cryo-EM) structures ^18–20^. The precise role of *N*-glycosylation in modulating ATG9A structure, function and dynamics remains unclear. The complex architecture and inherent flexibility of glycans hamper their experimental characterization using structural biology techniques ^23^. Molecular Dynamics (MD) simulations, however, can offer valuable insights into the conformational ensembles of *N-*glycans and glycoproteins ^24^. Prompted by these observations, we ran MD simulations of the ATG9A homotrimer in lipid bilayers to investigate the role of *N*-glycosylation at N99 on ATG9A protein dynamics and structure and disclose different mechanistic aspects. Additionally, we evaluated if the removal of the N-glycosylation of AT9A could alter its biological function at the autophagosome, using autophagy flux as a biological readout and glycosylation-null mutants of ATG9A (ATG9A^N99A^ and ATG9A^N99D^).

## Results and Discussion

### ATG9A undergoes asymmetric closed-open conformational changes in the non-glycosylated and glycosylated state

The ATG9A loop, which includes the N-glycosylated residue N99, is positioned at the interface between the solvent and the lipid headgroups of the membrane, close to the luminal opening of the protein’s central cavity (**Figure 1A**). This positioning prompted us to investigate whether *N*-glycosylation of N99 modulates the conformational states of ATG9A. To this end, we ran 1μs-long MD trajectories by conventional (deterministic) sampling of the homotrimeric ATG9A (residues 36-522) in both its N-glycosylated (ATG9A_N99glyco_) and non-glycosylated forms (ATG9A) (**Table 1**), embedded in a lipid bilayer composed of 1-palmitoyl-2-oleoyl-sn-glycerol-3-phosphocholine (POPC) and cholesterol (**Figure 1A-B**).

**Table 1.** Summary of the collected simulations.

| System | Force Field | Lipid bilayer composition | Total atoms | Lenght Time (μs) | N° replicates |
| --- | --- | --- | --- | --- | --- |
| **ATG9A** | CHARMM36m | POPC 75%<br>CHOL 25% | 323637 | 1 | 3 |
| **ATG9A<sub>N99glyco</sub>** | CHARMM36m | POPC 75%<br>CHOL 25% | 354144 | 1 | 3 |

Because a single unbiased MD trajectory is likely to sample only a limited portion of the conformational space, we concatenated the ATG9A and ATG9A_N99glyco_ trajectories into two 3μs macro-trajectories and analyzed these, along with the single replicates, to improve sampling and capture a broader range of conformational states.

Cryo-EM data show that ATG9A protomers (named protomer A, B and C) can adopt two distinct conformations, referred to as “open” and “closed” states in the regions surrounding the central cavity (**Figure 1C-D**)^18–20^. These conformations are defined by the degree of separation between the domain-swapped transmembrane helices (TMH3-4) and the rest of the same protomer as well as the TMH3–4 helices of the neighboring protomers at the interfaces (AC, BA, and CB, named after the interacting protomers, **Figure 1D**). In the open state, this separation is larger, while in the closed state, it is smaller.

We monitored two different parameters to characterize transitions between the closed and open states of ATG9A and ATG9A_N99glyco_, namely the Root Mean Square Deviation (RMSD) of the Cα atoms of the domain-swapped transmembrane helices of each protomer (TMH3-4) relative to the closed experimental cryo-EM structures (**Figure 1E** and **Figure S1-2**) and a contact score based on the occurrence of protein-protein contacts among the TMH3–4 helices of each protomer and other regions of ATG9A that are present only in the cryo-EM structure of the closed state (**Figure 1F** and **Figure S3**). We observed that in both ATG9A and ATG9A_N99glyco_, at least one protomer spontaneously transitioned towards the open state when starting from a closed state (**Figure 1E-F** and **Figure S1**).

^23, 25^The asymmetric open states of the protomers obtained from the MD simulations (**Figure 1E-F**) do not fully capture the complete open state observed in the cryo-EM data, which consistently shows a symmetric orientation of the protomers in all ATG9A structures^18,20^. This suggests the presence of a high energy barrier that prevents reaching the fully open state during the simulations. Further analysis of the cryo-EM data for ATG9A pointed out that the high-resolution structure determination in a nanodisc environment (PDB ID 7JLP, resolution 3.4 Å) ^20^ was based on approximately only 2.4% of the initial particle images. Similarly, the highest resolution structures of ATG9A in detergent for the open (PDB ID: 6WQZ, resolution 2.8 Å) and closed (PDB ID: 6WR4, resolution 2.9 Å) states ^18^ were derived from processing only 8.5% and 9.3% of the initial 2D particle images, respectively. These observations suggest that the available cryo-EM data may not fully rule out the existence of structural heterogeneity and additional conformational states beyond the reported open and closed configurations. This raises the possibility that certain states within the cryo-EM data could reflect the asymmetric conformations observed in the MD simulations.

The closed-to-open conformational changes in the MD trajectories are driven by the separation of the domain-swapped interfaces of the protomers (**Figure 1E-F**), resulting in a lateral opening of the central cavity towards the hydrophobic portion of the lipid bilayer. These data align with findings from cryo-EM data and from previous all-atom MD simulations ^20^. The opening of the domain-swapped interfaces involves interactions between the reentrant membrane helix 1 (RMH1)-turn-helix α9 of each protomer and TMH3–4 (**Figure 1C-D**). This behavior is evident in replicate 1 of ATG9A_N99glyco_, where all three domain-swapped interfaces shifted closer to the open state (**Figure S1** and **Figure S3**). Conversely, we also observed states in ATG9A_N99glyco_ with a high occurrence of specific contacts (i.e., contact score values close to 1), indicating that closed conformations are preserved in certain regions (**Figure 1F**). For example, this is reflected in the interface BA of replicate 2 and interface AC of replicate 3 in ATG9A_N99glyco_ (**Figure S3**). Additionally, both ATG9A and ATG9A_N99glyco_ can adopt conformations with high RMSD relative to both the experimental close and open structures. This is exemplified by the interface CB in replicate 2 of ATG9A_N99glyco_, (**Figure 1E and Figure S2**). These conformations provide alternative orientations of the TMH3–4 helices that are different from those observed in the final cryo-EM models ^18–20^.

### N-glycosylation at N99 favors the extent of open conformations in ATG9A

We performed clustering of trajectory frames based on contact scores at the domain-swapped protomer interfaces. We used Euclidean distance, while accounting for ATG9A’s threefold symmetry (i.e., interfaces AC, BA and CB), to gain deeper insights into the protein dynamics and identified diverse conformational states sampled by ATG9A and ATG9A_N99glyco_ (**Figure 2 and Figure S4**). A quality-threshold clustering approach^25^ (threshold 0.45) was applied, discarding clusters with fewer than 60 frames, and representative structures were identified based on minimal mean square distance within each cluster. In our MD simulations, cluster analysis and examination of representative structures unveiled different open and closed states in glycosylated ATG9A that are not at all or only rarely observed in non-glycosylated ATG9A and vice versa (**Figure 2A-B and Table S1**). To compare ATG9A and ATG9AN99glyco, we calculated Jaccard and weighted Jaccard similarity scores to quantify differences in contact score distributions (**Figure 2C and Table S1**). Further structural stratification was conducted using kernel density analysis of RMSD values for transmembrane TMH3-4 helices, referencing experimental cryo-EM structures of the open and closed states. ATG9A_N99glyco_ exhibited a higher occurrence of open conformations at all protomer interfaces compared to ATG9A. These open states were predominantly found in cluster 2 of ATG9A_N99glyco_, which encompassed approximately 30% of the concatenated trajectory frames. This cluster displayed representative contact scores of 0.2, 0.05, and 0.15 for interfaces AC, BA, and CB, respectively (**Figure 2B**). In contrast, ATG9A displayed only a partial population of such open states. Specifically, cluster 1 (comprising around 56% of the concatenated trajectory frames) (**Figure 2A**) and cluster 5 (representing around 2% of the frames) showed limited similarity with cluster 2 of ATG9A_N99glyco_, as indicated by the low Jaccard similarity values of approximately 15% and 10%, respectively (**Figure 2C and Table S1**). Cluster 1 of ATG9A included partially open and asymmetric states, with average contact scores of 0.35, 0.2, and 0.15 for the AC, BA, and CB interfaces, respectively, indicating a more closed conformation of the interfaces (**Figure 2A**). Similar asymmetric states, characterized by one or two open interfaces while the others remained closed, were observed in clusters 1, 3, 4, 6, and 7 of ATG9A_N99glyco_ and clusters 2, 3, and 4 of ATG9A. ATG9A_N99glyco_ exhibited a higher frequency of states involving at least one interface with a particularly high contact score (above 0.5), particularly in clusters 1 and 5 (**Figure 2B and Figure S4**). In contrast, these states were less common in ATG9A and appeared in clusters 2, 3, and 4, showing limited overlap with clusters 1 and 3 of ATG9A_N99glyco_ (**Figure 2C and Table S1**). The presence of these closed states at domain-swapped interfaces might be influenced by the initial structure or be intermediate states during closed-open conformational changes. We thus further analyzed the individual replicates by examining collective contact scores and the RMSD of domain-swapped interfaces relative to the experimental cryo-EM structures (**Figure S5**). While open conformational states were observed across replicates, there were distinct differences in their sampling behavior. Replicate 1 of ATG9A_N99glyco_ predominantly sampled open states for all three protomers, whereas replicate 2 and 3 primarily sampled conformations with two open interfaces and one closed interface (BA and AC, respectively). Further analysis of these states suggested that the ability of ATG9A_N99glyco_ to maintain closed domain-swapped interfaces could be linked to interactions between *N*-glycans and the luminal loops of neighboring protomers. Investigation of the surrounding environment of the three N*-*glycosylation revealed heterogeneous and transient interactions with glycan, protein, and lipid atoms (**Figure S6**). Moreover, in the second half of replicate 2, we observed an increased presence of protein and lipid atoms near the *N*-glycan of protomers C and B. These interactions may play a role in modulating the open or closed states of the domain-swapped interfaces.

**Figure 2.**
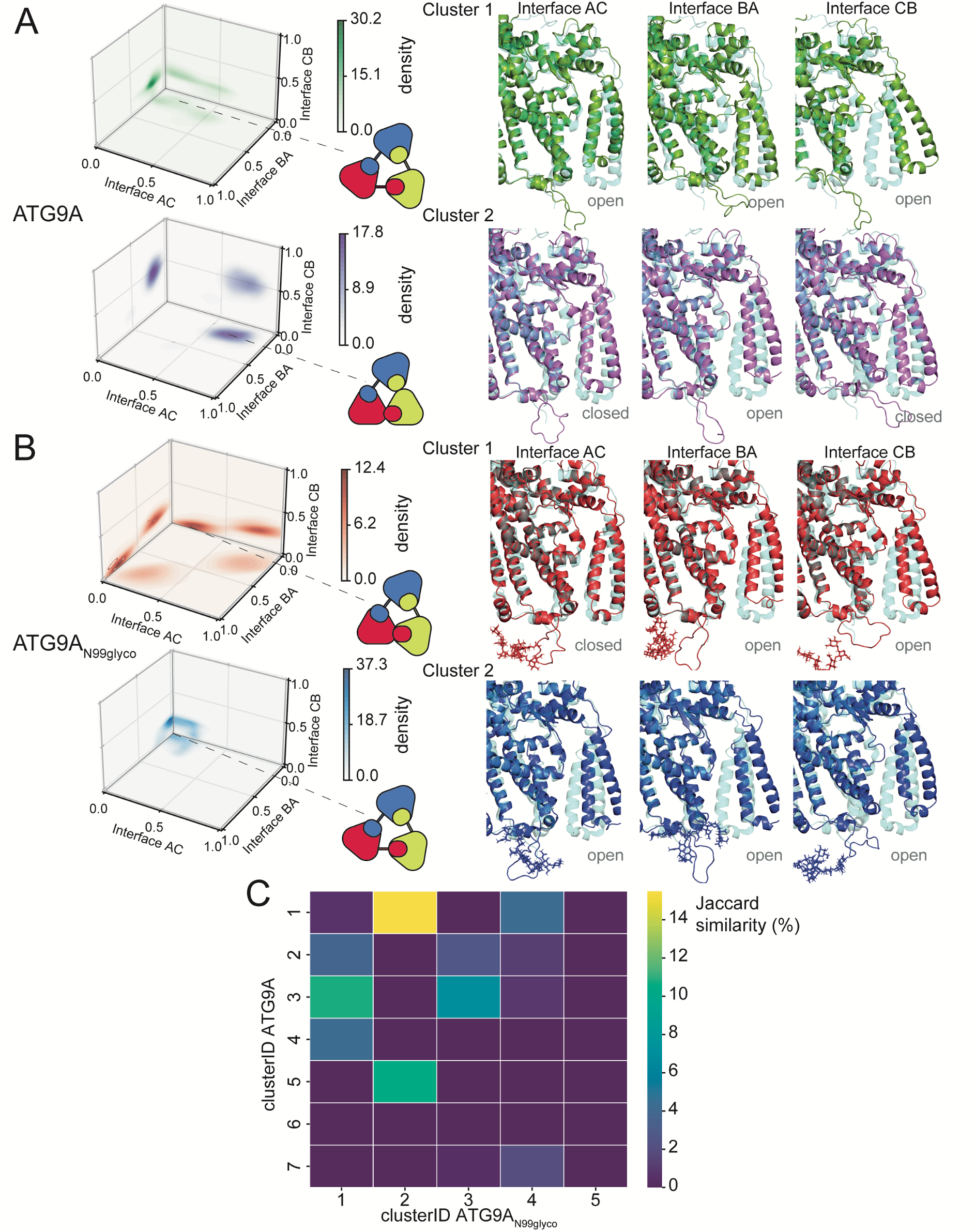
N-glycosylation at N99 modulates open-closed states of ATG9A homotrimer. The cluster analysis was conducted on the concatenated trajectories of ATG9A (A) and ATG9A_N99glyco_ (B). The analysis employed an Euclidean distance metric based on the contact scores at each protomer-protomer interface (interfaces AC, BA, and CB). A quality-threshold algorithm was employed with a set threshold of 0.45. The left panels display the bi-dimensional density projections of the contact score values for the two main clusters of ATG9A (A) and ATG9A_N99glyco_ (B). The right panels provide detailed highlights of the AC, BA and CB interfaces for the representative structures from the first and second main clusters identified. The structures are depicted using a cartoon model, and for reference purposes, they are superimposed on the closed-state structure, which is illustrated as a blue, semi-transparent cartoon. The N99 *N*-glycosylation are represented as sticks. (C) Heatmap of the Jaccard similarity index, calculated to evaluate the occurrence and commonality of states within each identified cluster for ATG9A and ATG9A_N99glyco_. A Jaccard similarity index value of 100% indicates identical state sets, while 0% indicates no similarity. ATG9A_N99glyco_ shows a higher occurrence than ATG9A of states with open conformations at all protomer interfaces (as seen in cluster 2, characterized by representative contact scores of 0.2, 0.05, and 0.15 for interfaces AC, BA, and CB). ATG9A only partially populates such open states (in cluster 1 and cluster 5), showing low similarity with cluster 2 of ATG9A_N99glyco_. ATG9A and ATG9AN99glyco in cluster 1 are asymmetric states, characterized by either two or one open interfaces, while the remaining interfaces are in a closed state.

Overall, our MD simulations suggested that the glycosylation of N99 favored the extent of open conformations. The occurrence of closed conformations in ATG9A_N99glyco_ associated with interactions between *N*-glycans from different protomers during the simulation should be taken cautiously, as inherent force-field limitations might influence them^24^.

### N99 glycosylation promotes cooperativity in the conformational changes of ATG9A

The analysis of the contact score presented above enabled us to discriminate between closed and open conformations of the individual ATG9A protomers relative to the cryo-EM reference states. Cooperativity in conformational changes is a hallmark of oligomeric proteins, playing a critical role in their functional modulation^26,27^. To investigate this aspect, we employed a statistical framework to explore the relationships among the open/closed states of each protomer (A, B, and C) of ATG9A and ATG9A_N99glyco_ with the open/closed state of the other protomers. Our goal was to evaluate if cooperativity played a role in the closed-to-open conformational change and in the effects of the N-glycosylation on this mechanism.

We used Log-Linear Models (LLMs) with the Poisson family to fit the contact score data and calculate deviance and log-likelihood (**Figure 3A and Figure S7**). These models are well-suited for modeling relationships among categorical variables, effectively capturing the complex interplay and dependencies between the protomer states. To evaluate the performances of different LLMs, we applied chi-squared statistics and calculate deviance and log-likelihood (**Figure S7**).

**Figure 3.**
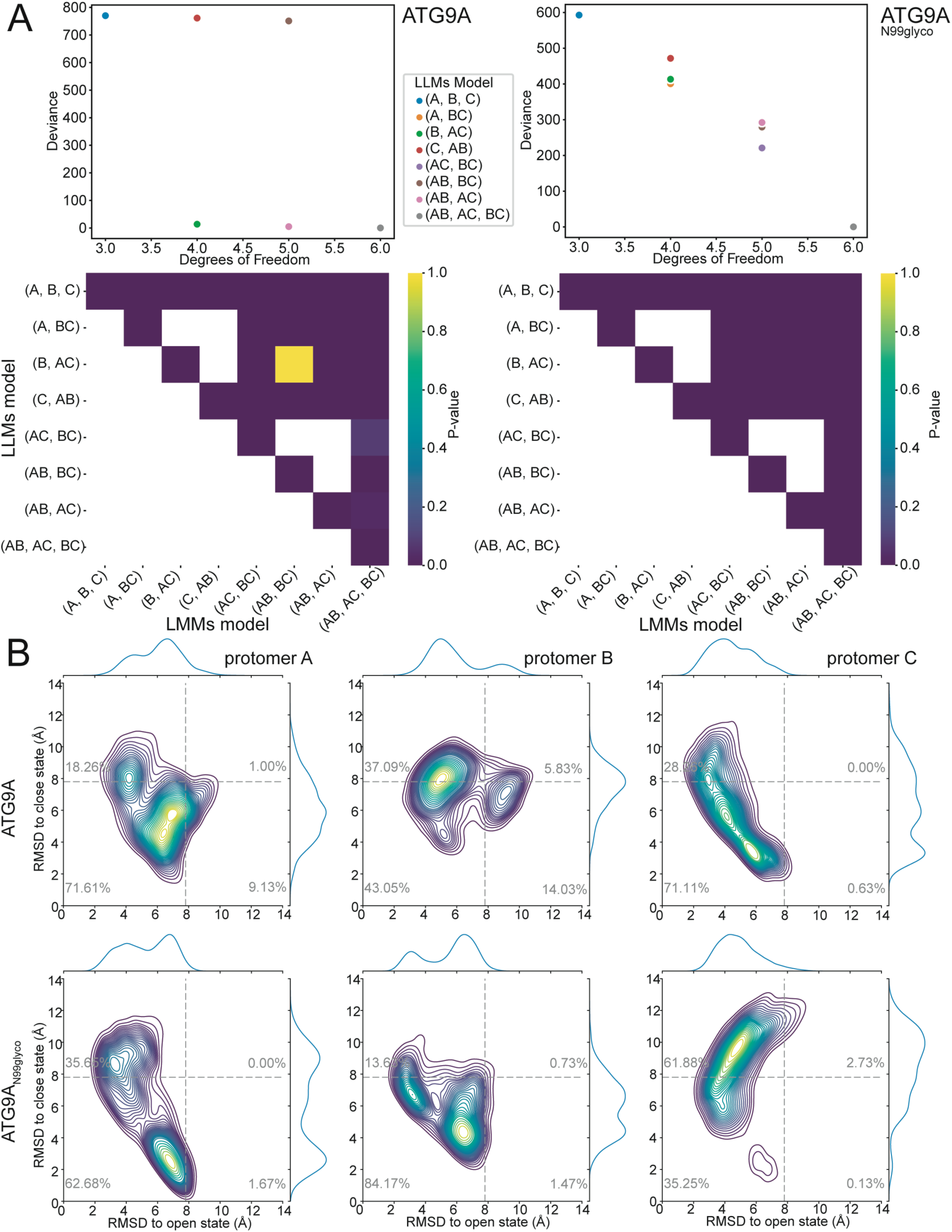
N-glycosylation promotes cooperativity in the closed-open conformational changes of ATG9A. (A) Log-Linear Models (LLMs) analysis illustrating the relationships among open/closed states of the protomers (A, B, C) of ATG9A (left panels) and ATG9A_N99glyco_ (right panels). The upper panels show the degrees of freedom for the different LLMs and their deviance in fitting the contact score data, highlighting differences in the fit of independence models between ATG9A and ATG9A_N99glyco_. We tested complete independence ([A, B, C]), joint dependence ([A, BC], [B, AC], [C, AB]), conditional independence ([AC, BC], [AB, BC], [AB, AC]) and homogeneous association model (i.e., [AB, AC, BC]). The lower panels show the p-value calculated as heatmaps to assess the significance of the deviance between different models. A low p-value (i.e., 0.05) suggests that the difference in deviance between the two compared models is statistically significant, i.e., unlikely to have occurred by chance. The homogeneous association model (i.e., [AB, AC, BC]) showed a good fit of the contact score data of ATG9A_N99glyco_, with the lowest deviance and highest log-likelihood (Figure S7) across all models, suggesting a dependent relationship amongst all three protomers, which is less apparent in ATG9A. (B) Kernel density distribution plots of the Root-Mean-Square Deviation (RMSD) values of domain-swapped regions for each protomer (A left panels, B middle panels, C right panels) relative to the cryo-EM closed and open structures. The analysis has been performed on the concatenated trajectories of ATG9A (upper panels) and ATG9A_N99glyco_ (lower panels). The dotted grey lines indicate the 7.8 Å threshold for RMSD values, calculated from comparing open and closed cryo-EM structures. ATG9A exhibited a higher occurrence of alternative states with high RMSD values relative to both open and closed experimental structures (1%, 6%, 0% for interfaces AC, BA, and CB, respectively), compared to ATG9A_N99glyco_ (0%, 0%, 2%).

For both ATG9A and ATG9A_N99glyco_, the complete independence model ([A, B, C]) exhibited higher deviance and lower log-likelihood, indicating a poor fit and suggesting dependencies in the open/closed states among the protomers (**Figure 3A and Figure S7**). In ATG9A, both the homogeneous association model ([AB, AC, BC]) and conditional independence models, particularly [AC, BC] and [AB, AC], resulted in low deviances and high log-likelihoods. These findings suggest potential dependencies among the open/closed states of all protomers, or a scenario where the status of two protomers (A, B and B, C) is influenced by the status of the third (C and A, respectively).

Comparatively, ATG9A_N99glyco_ displayed a distinct relationship in protomer open/closed states. The joint dependence ([A, BC], [B, AC], [C, AB]), and conditional independence models did not show a better fit than the complete independence model (**Figure 3A and Figure S7**). Notably, for ATG9A_N99glyco_ the homogeneous association model ([AB, AC, BC]) provided the best fit for the contact score data, with the lowest deviance and highest log-likelihood among all models. This suggests a dependent relationship among all three protomers, with their open/closed states collectively influenced—an effect less apparent in ATG9A. Overall, our results provide a model in which glycosylation at N99 enhances cooperative interactions among the protomers, potentially modulating the closed-open conformational changes.

Building on these results, we further analyzed the open and closed states of individual protomers to evaluate the existence of alternative conformations not captured in the deposited cryo-EM structures. To this end, we performed RMSD calculations for the domain-swapped regions of each protomer relative to the experimental cryo-EM structures (**Figure 3B**). The classification of protomer states was based on structural comparisons. If the RMSD between a given protomer conformation and the reference open or closed structure exceeded 7.8 Å, it was considered a distinct state. This threshold was chosen based on observed differences between open and closed cryo-EM structures. ATG9A_N99glyco_ exhibited a higher occurrence of open-like states at the protomer interfaces (36%, 14%, 62% for interfaces AC, BA, and CB, respectively) compared to ATG9A (18%, 37%, and 28%) (**Figure 3B**). Conversely, ATG9A showed a higher occurrence of alternative states with high RMSD values relative to both open and closed experimental structures (1%, 6%, 0% for interfaces AC, BA, and CB, respectively) than ATG9A_N99glyco_ (0%, 0%, and 2%). This difference became more pronounced for ATG9A when considering states with RMSD values in the 6-8 Å range (**Figure 3B**). These findings suggest that *N*-glycosylation at N99 may contribute to restricting the conformational flexibility of ATG9A, limiting the occurrence of alternative states that deviate from the known open and closed experimental structures.

### Conformational changes modulate the shape and lipid accessibility of the central cavity of ATG9A

The different conformational states observed in ATG9A and ATG9A_N99glyco_ may influence the protein internal cavities and their role in modulating lipid movements. To investigate these differences, we analyzed the internal cavities of ATG9A, specifically focusing on variations in cavity profiles between the open and closed states. Our analysis targeted the shape (i.e., radius and length) and accessibility of the central cavity formed by the three protomers (**Figure 4**), as this cavity is proposed to mediate lipid scrambling between membrane leaflets^18–20^. We carried out a similar analysis on the cryo-EM-derived models of the open and closed states for reference (**Figure 4A**). Consistent with previous findings^18,20^, our analysis of the two reference structures revealed that the central pore is predominantly hydrophilic, highly solvated and spans approximately 60 Å.

**Figure 4.**
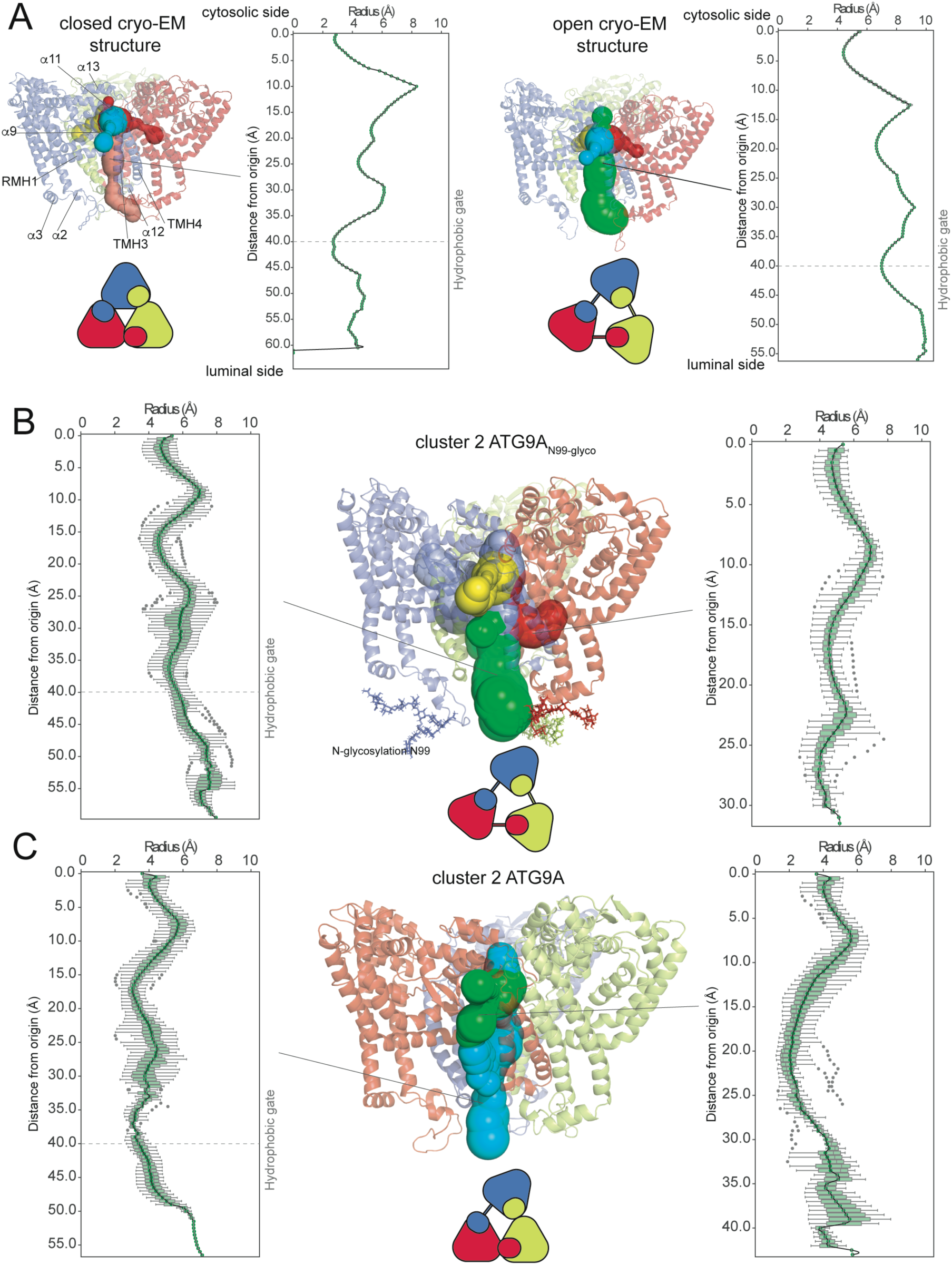
Closed-open conformational changes of ATG9A modulate the radius and solvent accessibility of the central cavity of ATG9A. Analysis using CAVER of the radius and length of the central cavity and its side cavities. (A) Analysis of the cavities of the cryo-EM-derived models of the closed (left panels) and open (right panels) states. (B-C) Analysis of the cavities of cluster 2 of ATG9A_N99glyco_ (B) and ATG9A (C). The box plots show the average profiles of radius and length (calculated as the distance from the starting point for the cavity calculation) for the central cavities calculated across all the structures within each cluster. The ATG9A structure is shown as a cartoon, highlighting its three protomers: A (red), B (blue), and C (yellow). The cavities identified are represented as spheres colored based on the tunnel clusters. Each panel includes a schematic representation showing the open and closed asymmetric conformations of ATG9A at the protomer interfaces. The average profile of the central cavity from cluster 2 of ATG9A_N99glyco_ is similar to the reference open state, showing shorter cavities that open sideways at each domain-swapped interface towards the hydrophobic environment of the bilayer.

The central cavity is constrained at the cytosolic entrance by helices α11 and α13 and, at the luminal side, by a hydrophobic gate at its narrowest point (cavity radius ∼2.5 Å in the closed state) formed by TMH3-4 and helix α12. The luminal opening of the central cavity is further modulated by the luminal loop—which includes the *N*-glycosylation site— and the helices α2 and α3 (**Figure 4A**). The main difference between the open and closed states of ATG9A lies in the increased degree of cavity opening in the luminal region with the hydrophobic gate radius expanding to ∼5.5 Å in the open state (**Figure 4A**). In both the open and closed structures, we identified three groups of side cavities adjacent to the central cavity. Each group corresponds to a domain-swapped interface and extends beneath α11 on the cytosolic side of the protein (**Figure 4A**). Lipid molecules were observed inserting their headgroups into these cavities in one of the available cryo-EM structures^20^.

We further examined the central cavity and each cluster in ATG9A and ATG9A_N99glyco_ (**Figure 4**) from our cluster analysis. We calculated average profiles of the radius and length of the cavities across all frames within each cluster (**Figure 4B-C**). In the open states sampled by ATG9A_N99glyco_, particularly cluster 2 (encompassing ∼30% of the frames), the central cavity exhibited an average radius and length profile closely matching the reference open state (**Figure 4B**). Additionally, our analysis identified shorter cavities branching sideways from the central cavity at each domain-swapped interface, extending beneath RMH1-turn-helix α9 and TMH3–4 toward the hydrophobic bilayer environment (**Figure 4B**). In contrast, for ATG9A, cluster 2 states with two closed interfaces (AC and CB), showed an average central cavity profile resembling the reference closed state with a radius of ∼ 3.5 Å around the hydrophobic gate (**Figure 4C**). In addition, the central cavity at the interface BA, which adopted open states, exhibited a sidewise opening (**Figure 4C**, right panel).

In the proposed model for ATG9A scrambling activity, the central cavity facilitates the trans-bilayer movement of phospholipids, with lipid molecules inserting into the cavity in both cryo-EM studies and MD simulations^20^. To further investigate the role of the N99 glycosylation in this mechanism, we monitored the entrance of phospholipids into the central cavity of ATG9A and their subsequent trans-bilayer movements.

We analyzed the evolution of the shape of the bilayer surface during the simulations and the lipid headgroup positions and their interactions with the protein using LipidDyn^28^. Lipids that reduced their distance to the opposite leaflet by at least 20% compared to the reference distance between the cytosolic and luminal leaflets were classified as undergoing partial translocation across the bilayer (**Figure 5A**). In ATG9A_N99glyco_, a higher average of phospholipids exhibited trans-bilayer movements compared to ATG9A (34 versus 27 phospholipids, respectively) (**Figure 5A**). This suggests that the N99 glycosylation in ATG9A could facilitate partial trans-bilayer movements of phospholipids. Additionally, we investigated cholesterol translocation and observed that cholesterol molecules underwent trans-bilayer movements almost five times more frequently than phospholipids. However, no differences were detected between ATG9A and ATG9A_N99glyco_ in terms of cholesterol translocation (**Table S2**).

**Figure 5.**
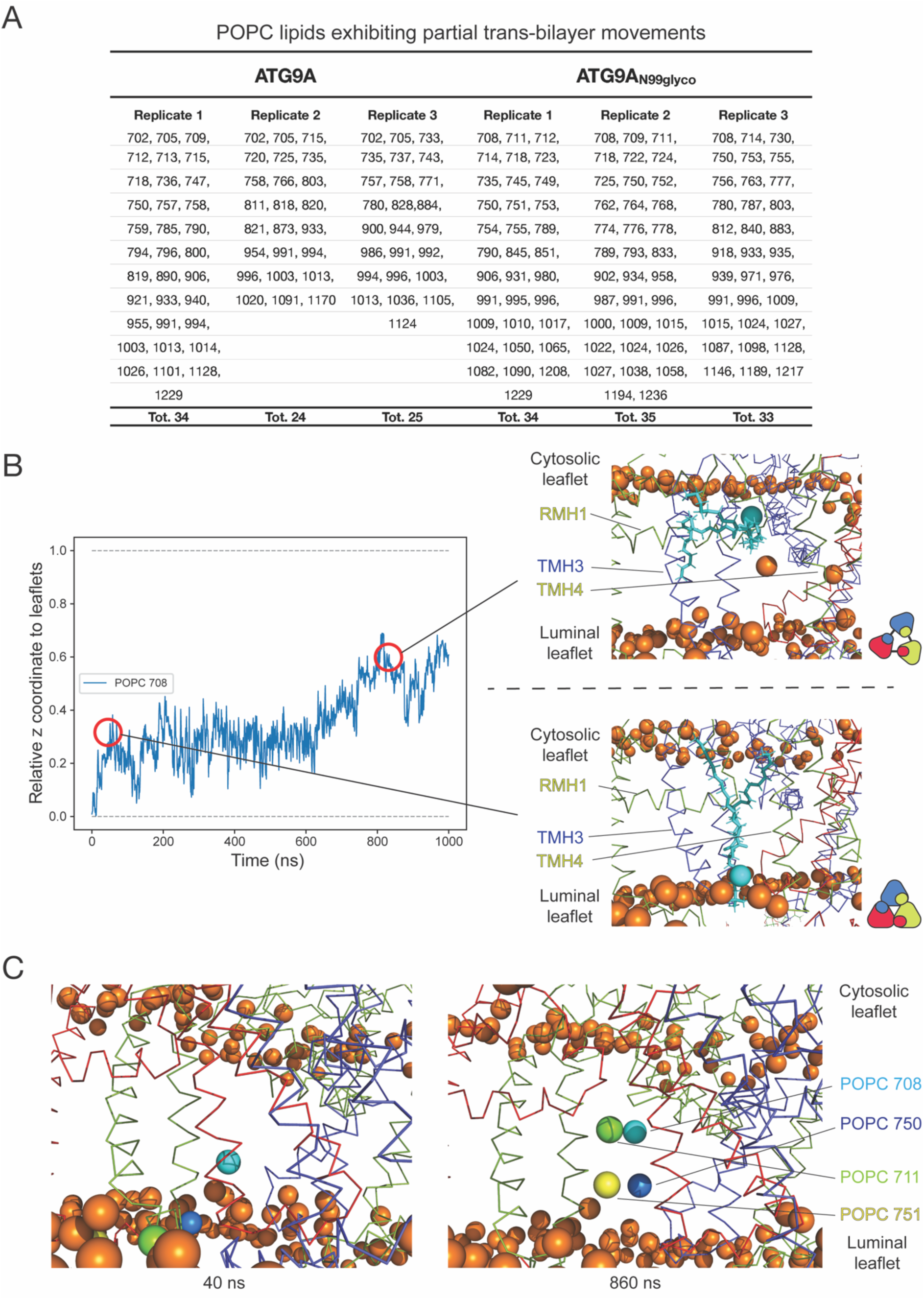
ATG9A N-glycosylation favors open states of the central cavity and facilitates partial trans-bilayer movements of phospholipids. (A) Analysis using LipidDyn of the POPC lipids showing partial trans-bilayer movements for each replica of ATG9A and ATG9A_N99glyco_. In ATG9A_N99glyco_, we observed a higher average of phospholipids exhibiting partial trans-bilayer movements compared to ATG9A. (B) Example of a lipid (POPC 708) inserting into the central cavity of ATG9A and undergoing reorientation and partial trans-bilayer movement in replicate1 of ATG9A_N99glyco_. The left panel shows the time-wise relative position along the z axis of the head group of POPC 708 (blue line) with respect to the two leaflets. Here the value of 1 corresponds to the cytosolic leaflet and 0 corresponds to the luminal leaflet. The right panels show two representative frames at 30ns (lower) and 860ns (upper) of the simulation. The ATG9A structure is shown as a ribbon, highlighting its three protomers: A (red), B (blue), and C (yellow). Lipids headgroups are shown as orange spheres while POPC 708 is highlighted in cyan. The lipid headgroup is inserted in the central cavity at the domain-swapped interface CB (30 ns, lower) and then reorient horizontally while still maintaining the acyl tails in the favorable hydrophobic lipids’ bilayer (860ns, upper). Each panel includes a schematic representation showing the open and closed asymmetric conformations of ATG9A at the protomer interfaces. (C) Frame at 40ns (left panel) and frame at 860ns (right panel) extracted from replicate1 of ATG9A_N99glyco_. The headgroups of some of the lipids undergoing partial trans-bilayer motions, POPC 708 (cyan), POPC 711 (green), POPC 750 (blue) and POPC 751 (yellow), are shown as spheres. We observed that multiple phospholipids inserted inside the central cavity.

Our simulations revealed that phospholipids from the luminal leaflet inserted their hydrophilic headgroups into the protein central cavity at the domain-swapped interfaces, orienting horizontally while their acyl tails remained within the bilayer hydrophobic environment (**Figure 5B**). Multiple phospholipids inserted at the same time into the central cavity and exhibited partial trans-bilayer motions (**Figure 5C**). Specifically, phospholipids from the luminal leaflet traversed ∼60% of the distance between the luminal and cytosolic leaflets, reaching the region of RMH1 and TMH3–4 in each protomer (**Figure 5B-C**). Although additional simulations with enhanced sampling approaches would be necessary to observe complete trans-bilayer lipid movements, our findings suggest that this region may act as a gate or exit point for lipid movements, influenced by bilayer curvature.

In conclusion, our computational analyses point out a mechanism in which closed-to-open conformational changes resulted in sidewise openings and increased accessibility of the central cavity towards the hydrophobic region of the lipid bilayer. Additionally, the hydrophilic nature of the central cavity provides a favorable environment for the polar headgroups of phospholipids (**Figure 5B-C**). Open conformations of the protein facilitate lipid insertion into the central cavity, along with their reorientation and partial transbilayer movements, representing potential steps in the lipid scrambling mechanism. Furthermore, the wider radius of the central cavity in the open state (**Figure 4B**) allows it to accommodate the headgroups of diverse phospholipids, aligning with findings from scrambling assays^20^. N99 glycosylation overall facilitates the trans-bilayer movements.

### Effects of N-glycosylation of ATG9A on the autophagy flux

Based on the effects of *N*-glycosylation on the conformational changes of ATG9A and lipid movements through the protein cavities we determined by MD simulations, we investigated if these findings do translate into observable changes in the function of the protein and more specifically in its involvement in autophagy. We selected the analysis of the autophagy flux as biological readout for cellular experiments^29^. Autophagic flux can be evaluated by measuring the turnover of key autophagy markers like LC3-II and SQSTM1/p62 under conditions that block lysosomal degradation, such as treatment with Bafilomycin A1^30,31^. This approach distinguishes between autophagosome synthesis and degradation phases. In our case, steady-state measurements alone are insufficient, as they may not capture alterations induced by specific genetic modifications, such as ATG9A variants, thus it is necessary to normalize the measurements using specific ratios to accurately quantify autophagy flux ^32^.

To assess the effects of ATG9A *N*-glycosylation on autophagy flux, we generated two ATG9A variants, ATG9A^N99A^ and ATG9A^N99D^, which deny the *N-*glycosylation sequon and thus impair *N*-glycosylation at the site. To assess the effects of ATG9A N-glycosylation on autophagy flux, we first investigated the potential pathogenicity of variants at the N99 and S101 sites, as both can impact the N-glycosylation sequon. We examined available data from gnomAD^33^ and ClinVar^34^ and found no reported pathogenic or likely pathogenic variants at either site. In contrast, Alphamissense^35^ predicts several N99 variants (N99W, N99V, N99M, N99I, N99F, and N99C) as likely pathogenic, while no pathogenic nor likely pathogenic variants were reported for S101. Based on these findings, we generated two ATG9A variants, ATG9A^N99A^ and ATG9A^N99D^, which deny the *N-*glycosylation sequon and thus impair *N*-glycosylation at the site. To ensure that the endogenous ATG9A levels did not interfere with the analysis, we used both ATG9A knockout (KO) and control HEK293 cell lines. The ATG9A KO cell line provided a clean background where only the introduced ATG9A variants were expressed, allowing us to specifically evaluate their role in autophagy flux without the confounding effects of endogenous ATG9A. Autophagy flux was assessed by comparing LC3-II and SQSTM1 turnover in the presence and absence of bafilomycin A1, a lysosomal degradation inhibitor. This approach enabled us to determine whether the ATG9A variants enhance or impair autophagy by analyzing both markers synthesis and degradation ratios. First, we validated the KO cell line by confirming that no levels of endogenous ATG9A were detected (**Figure 6A**), then, upon transient ATG9A variant overexpression, we measured the variants’ effects on the autophagy marker synthesis and degradation ratios as compared to ATG9A WT (**Figure 6**). Our results did not show any statistically significant impact of the ATG9A^N99A^ and ATG9A^N99D^ variants compared to ATG9A WT (**Figure 6A-C**). This conclusion was based on the analysis of both LC3-II and SQSTM1/p62 levels, which showed no significant differences under the tested conditions.

**Figure 6.**
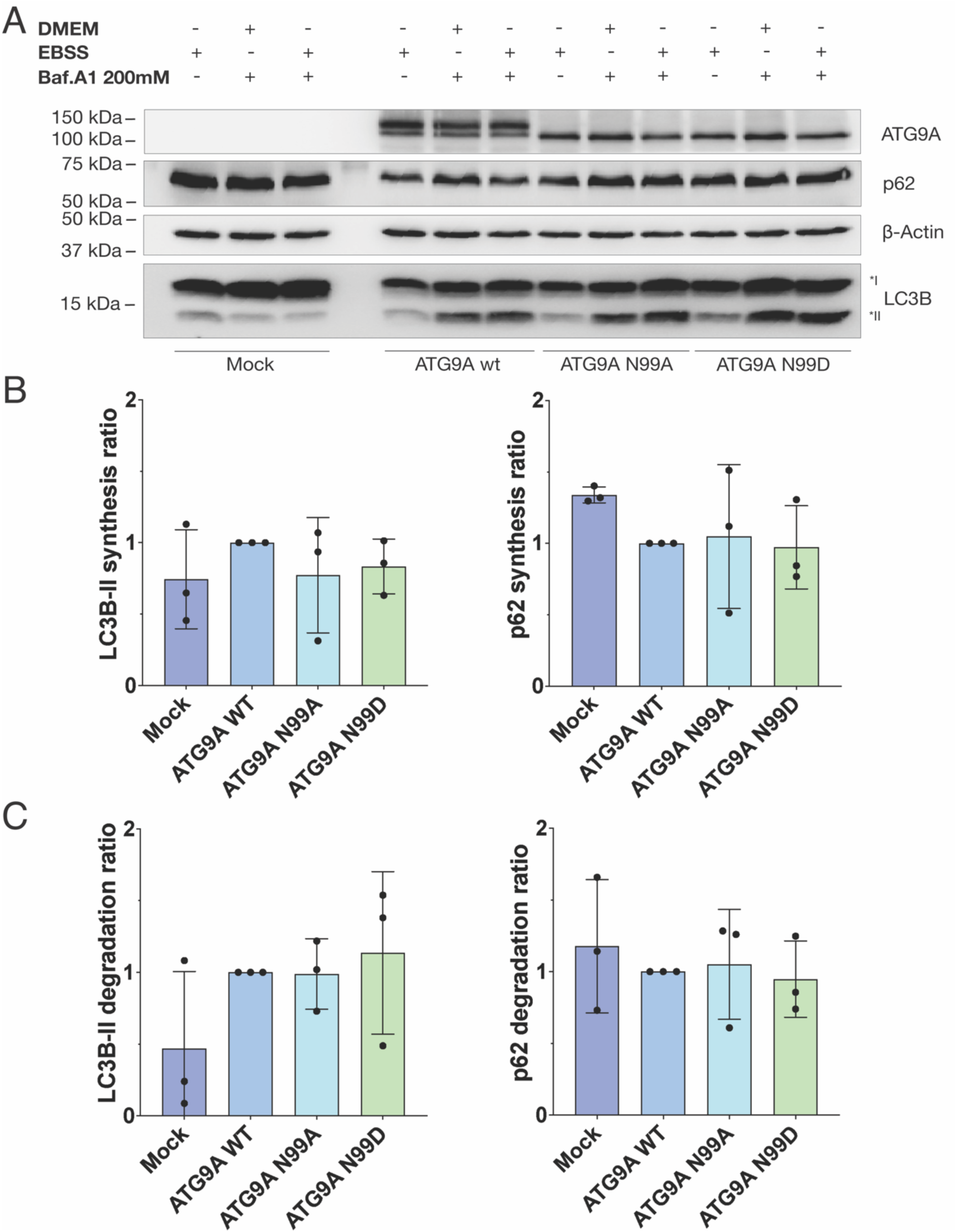
Abolishing *N*-glycosylation through mutagenesis is not sufficient to capture any difference in autophagosome synthesis and degradation in the ATG9A KO HEK293 cell line. (A) Western blot analysis of the LC3-II and p62 levels of ATG9A WT, ATG9A^N99A^ and ATG9A^N99D^ under starvation (EBSS medium) or rich-nutrient conditions (DMEM medium), in the presence of bafilomycin A1, a lysosomal degradation inhibitor, in the ATG9A KO HEK293 cell line. ATG9A^N99A^ and ATG9A^N99D^ samples were both loaded on a 15% gel along with ATG9A WT and the Mock condition (i.e., no overexpression). LC3-II and SQSTM1/p62 levels have been normalized with the housekeeping protein β-Actin levels in the same sample. (B) Analysis of the autophagosome formation rate (synthesis rate) following LC3-II (left) or SQSTM1/p62 (right) levels. The synthesis rate has been calculated as the rate of LC3-II (or SQSTM1/p62) levels in the presence of the inducer (i.e., EBSS) and the inhibitor (i.e., Bafilomycin A1) divided by the LC3-II (or SQSTM1/p62) level in the presence of the inhibitor alone. (C) Analysis of the autophagosome degradation rate following LC3-II (left) or SQSTM1/p62 (right) levels. The degradation rate has been calculated as the rate of LC3-II (or SQSTM1/p62) levels in the presence of the inducer (i.e., EBSS) and the inhibitor (i.e., Bafilomycin A1) divided by the LC3-II (or SQSTM1/p62) level in the presence of the inducer alone.

## Materials and Methods

### Preparation of the starting structure

We used as starting structures for the MD simulations the cryo-EM structures of human ATG9A homotrimer in the closed state (ATG9A, PDB entry 7JLP chain A, B, and C, ^20^) and designed a construct that includes the residues 36-522. We selected this structure of ATG9A since it is the only one so far obtained in nanodiscs (resolution of 3.4 Å), using lipid composition of 1-palmitoyl-2-oleoyl-sn-glycero-3-phosphocholine (POPC): 1-Palmitoyl-2-oleoyl-sn-glycero-3-phosphatidylserine (POPS) at a 3:1 ratio. In the structure of ATG9A, we removed the residues 523-532 that are not solved in other structures of ATG9A and formed a disordered region on top of the cytosolic side of ATG9A. We removed the lipid molecules that were present in the cryo-EM structures. We used MODELLER v. 9.15 ^36^ to reconstruct the missing coordinates in the region 96-108, keeping the rest of the coordinates in the original positions. We generated 100 different models with MODELLER for ATG9A. We then used the statistically optimized atomic potential protein orientation-dependent score (SOAP-Protein-OD, ^37^) to evaluate the quality of the models and to rank them. SOAP-Protein-OD is an orientation-dependent potential. It is optimized for scoring protein models by evaluating the environment of residue in the models with respect to the expected environment from experimental structures. We used SOAP potential since it was developed to assess protein-protein interfaces and multiple-chain systems^37^. We also calculated the normalized DOPE z score and checked that the models do not have scores with positive values, which indicates low-quality models. We selected the top five models ranked on the SOAP-Protein-OD score, and we analyzed them by calculating Van der Waals clashes using Arpeggio^38^ and by visual inspection of the reconstructed regions. Between these five models, we selected one model for ATG9A in which the reconstructed regions have similar orientations between the protomers and do not make extensive contact with the rest of the protein to avoid a bias in the initial conformation for MD simulations. We used the selected model of ATG9A to design a glycosylated form of ATG9A (ATG9A_N99glyco_). ATG9A is glycosylated with a complex-type *N*-glycan at N99^4^, located in a loop facing the luminal side of the membrane. We modeled the *N*-glycosylation of each protomer of ATG9A_N99glyco_ with a common complex-type structure of *N*-glycans in humans^24^ using the tools available in CHARMM-GUI version 3.8^39^. *N*-glycans at N99 are complex, biantennary, mono sialylated and core fucosylated (GlyTouCanID G65761WK). This type of *N*-glycosylation is compatible with the localization of the protein in the trans Golgi.

### Design of ATG9A and ATG9A_N99glyco_ systems with lipid bilayers

We used the selected model of ATG9A and ATG9A_N99glyco_ to build the systems with lipid bilayers. We generated the systems by CHARMM-GUI version 3.6^39–41^. In CHARMM-GUI, we solvated the cavities of ATG9A. We used PyMOL to include in our systems only the water molecules inside the cavities using a cutoff of 4.5 Å for the distance between the water molecules and protein atoms lining the cavities. We identified the protein atoms lining the cavities of ATG9A by using CAVER 3.02 software^42^. We used as an initial starting point for the cavity calculations the so-called “center of gravity”^42^ defined by V440 of protomers A, B, and C. This residue is located on the top of the structure on the cytosolic side and oriented toward the central cavity of ATG9A. We used as parameters for the CAVER calculations a probe radius of 1.2 Å, comparable to the size of a water molecule^43^, a shell radius of 15 Å, a shell depth of 4 Å, and a clustering threshold of 8 Å. We included 307 water molecules inside the cavities of ATG9A. We included the same number of water molecules inside the cavities of ATG9A_N99glyco_.

We defined the protonation of the histidines by visual inspection of the experimental structures and by the predictions of PROPKA 3 ^44^ and H++ webserver^45^. We defined all the histidines as ε tautomer apart from H161 as δ tautomer. We capped the N- and C-terminus with the acetyl (ACE) and *N*-methylamide (CT3) groups. We inserted the protein inside a lipid bilayer with the size of 160, and 160 Å in the x and y dimensions, respectively, and composed of 1-palmitoyl-2-oleoyl-sn-glycerol-3-phosphocholine (POPC) 75% and cholesterol 25%. We here used a standard lipid composition for the bilayer since the detailed compositions of phagophore and autophagic membranes remain poorly understood due to challenges in isolating and characterizing these transient and dynamic structures^46,47^. The lipid bilayer was symmetric and included 717 lipids. To orient the protein in the membrane, we used the Orientation of Proteins in Membranes (OPM) webserver (https://opm.phar.umich.edu/ppm_server,^48^) in CHARMM-GUI. The OPM database uses the Positioning of Proteins in Membrane approach to determine the orientation of the protein in a membrane bilayer by minimizing its transfer energy with respect to several geometric variables in a coordinate system whose z axis coincides with the membrane bilayer normal. The systems were solvated in a rectangular box of water molecules with a minimum distance of 20 Å between the protein atoms and the edge of the water layers, including around 100 water molecules per lipid. We used a solvent model TIP3P adjusted for CHARMM force fields with Lennard-Jones sites on the hydrogen atoms^49^.

### Molecular Dynamics simulations

We equilibrated the systems according to the CHARMM GUI procedure that involves minimization followed by steps in which positional restraints for lipid head groups and protein backbone atoms are gradually removed. We carried out three replicates of 1-μs all-atom explicit solvent MD simulations for ATG9A and ATG9A_N99glyco_ with the CHARMM36m force field^50,51^ (**Table 1**). We performed the MD simulations with GROMACS^52^. We initialized each replicate using different atomic velocities. We carried out the simulations at a concentration of 0.15 M of NaCl, neutralizing the net charge of the system. We simulated each system under periodic boundary conditions. We used the LINCS algorithm to constrain the hydrogen bonds^53^ and apply a time step of 2 fs. We applied the particle-mesh Ewald (PME) summation scheme^54^ to calculate long-range electrostatic interactions using a Fourier spacing of 1.2 Å and interpolation order of 4. Van der Waals and Coulomb interactions were truncated at 12 Å. We performed productive MD simulations in the NPT ensemble at 310 K and 1 atm. We used a Nose-Hoover thermostat^55^ for temperature control, with coupling every 1 ps, and a Parrinello-Rahman barostat^56^ for pressure control, with coupling every 5 ps. We used semi-isotropic coupling in which the lateral pressure Pxy and normal pressure Pz dimension are coupled independently. Other details are provided in the input files in OSF (https://osf.io/c4aqu/).

We evaluated the evolution of the systems during the MD trajectories by calculating the Cα-atoms Root Mean Square Deviation (RMSD) and radius of gyration. We calculated the minimum distance between the protein atoms and its periodic images to find potential periodic boundary artifacts. In all the simulations, we observe a minimal distance always higher than 28 Å. We joined the three individual 1-μs replicates of ATG9A and ATG9A_N99glyco_ to obtain two 3-μs ensembles of conformational states (i.e. concatenated trajectories) that were used in the analysis.

### Analysis of conformational changes and protein flexibility

We calculated the RMSD values of all the Cα atoms with respect to the cryo-EM structures of the closed (PDB ID 7JLP)^20^ and open state (PDB ID 6WQZ)^18^. We excluded from the analysis the flexible loop (residues 96-111) that we reconstructed by MODELLER in the starting models. To better visualize the open/closed conformations that could potentially be sampled independently by each protomer, we then calculated i) RMSD of all Cα atoms of every single protomer (i.e., chain A, B, C) and ii) RMSD of the Cα atoms of only the swapped transmembrane helices of each protomer (named TMH3 and TMH4)^20^ with respect to the experimental cryo-EM structures of the closed and open state. In both cases, we used all the helical transmembrane regions of the protomers for the rigid body superposition (residues 43-55, 59-86, 119-127, 130-168, 180-193, 206-225, 245-256, 279-322 and 472-507 of each protomer).

We calculated a contact score based on analysis of the fraction of contacts present only in the experimental cryo-EM structure of the closed state when compared to the open one. We employed the software CONtact Analysis (CONAN)^57^ to analyze intramolecular contacts in the cryo-EM structures of the closed (PDB ID 7JLP) and open state (PDB ID 6WQZ) of ATG9A. We used a cutoff *r_cut_* value of 10 Å, *r_inter_* and *r_high-inter_* values of 5 Å, using a protocol previously employed^58^. We processed the data to obtain a list of contacts identified only in the closed structure. We then filtered these contacts, including only the ones formed by residues in the regions of ATG9A involved in the conformational changes. Thus, we included contacts between residues 367-431 of the exchanged (i.e., domain-swapped) transmembrane helices TMH3-4 of each protomer and region 316-335 of the same protomer, and region 389-426 of the corresponding swapping protomer. The domain-swapping order of protomers is i) protomer A with protomer C, ii) protomer B with protomer A, and iii) protomer C with protomer B. We then monitored the presence of these contacts during the MD trajectory and calculated a timewise contact score as the fraction of contacts, i.e., values close to 1 mean that nearly all the contacts are present and the structure is similar to the closed one, while a value close to 0 means that nearly all the contacts are absent. We calculated the per-residue Cα-atoms Root Mean Square Fluctuations (RMSF) as the flexibility index. We calculated RMSF as averaged over consecutive and non-overlapping time windows of 10 and 100 ns along the trajectories, as used for other proteins^59,60^. The analysis of the glycans, protein and lipid atoms in the surround of the N-glycans has been performed using a 6 Å distance cutoff.

### Analysis of conformational states and clustering

We further analyzed the results of the contact score to perform clustering of the trajectory frames. The distance between frames was determined by the Euclidean distance using the contact score values of each domain-swapped protomer interface as coordinates. To consider the threefold symmetry of the ATG9A homotrimer, we performed the clustering using the minimum of the distances to each of the three possible rotations of the protein for each pair of frames (e.g. a structure with one open interface AC and two closed interfaces BA and CB is considered equivalent to a structure with one open interface BA and two closed interfaces CB and AC). We implemented a script (see OSF, https://osf.io/c4aqu/) to perform quality-threshold clustering^25^ with threshold 0.45, discarding clusters with less than 60 frames (less 2% of the total). For each cluster, we determined its occurrence in the combined trajectories and its representative structure as the trajectory frame of the cluster with the minimum mean square distance to the rest of the frames in the cluster. To estimate the similarity between the sets of AC/BA/CB contact scores defined by each cluster in ATG9A and ATG9A_N99glyco_, we calculated as measurement the i) Jaccard similarity and ii) weighted Jaccard similarity to also take into account the frequencies of the values. The results are in both cases a value between 0 and 100%, where 100% indicates identical sets, and 0 indicates no similarity. To further stratify these sets, we calculated their associated kernel density distribution of the RMSD values of the Cα atoms of the swapped transmembrane helices TMH3-4 of each with respect to the experimental cryo-EM structures of the closed and open state.

### Independence analysis of the closed/open status of protomers

We used an in-house Python tool to evaluate the dependence/independence relationship of the open/closed status of each individual protomer of ATG9A (A, B, and C) with respect to the open/closed status of the other protomers (https://osf.io/c4aqu/). This allowed us to shed light on potential ongoing cooperativity in the conformational changes of the protomers. As input data for the analysis, we used the contact score values calculated for each protomer on the combined trajectories of ATG9A and ATG9A_N99glyco_. We applied a cutoff of 0.5 to classify the states as open or closed based on the distribution of contact score data, as this value appeared to effectively distinguish between them. As statistical models, we utilized Log-Linear Models (LLMs) with the Poisson family to fit the data and calculate deviance and log likelihood. These models are suitable for capturing the complex interactions and dependencies within the system and model the relationship between categorical variables (open/closed status of proteins) that have binary outcomes (open or closed) while considering potential nonlinear dependencies. The models investigated include complete independence, joint independence, conditional independence, and a homogeneous association model, each focusing on different combinations of relationships between the protomer open-closed status. We used chi-squared statistics to assess the difference in the deviance between different models and assess their goodness-of-fit and determine whether they are significantly different from each other. Furthermore, we employed Tukey Honestly Significant Difference (HSD) tests to analyze the deviance and log likelihood data, providing insights into the statistical significance of the model comparisons.

### Analysis of protein tunnels and channels

We used the standalone version of CAVER 3.02 software^42,43^ to monitor the cavities of ATG9A and how they evolve during the MD trajectories.

Due to the complex branched network of tunnels in ATG9A, we selected different starting points as a reference for searching cavities using CAVER. In detail, we selected: i) the centroid defined by considering all the atoms of K363 and K359 of all protomers (i.e., chain A, B, C) to monitor the central cavity, ii) the centroid defined by considering all the atoms of E323, A329, Y358 of a protomer and R422 of the corresponding swapped protomer (i.e., for chain A, we selected E323, A329, Y358 of chain A and R422 of chain C) to monitor each of the three lateral cavities, iii) the centroid defined by considering all the atoms of Y316, L354 of a protomer and F382 of the corresponding swapped protomer (i.e., for chain A, we selected Y316, L354 of chain A and F382 of chain C) to monitor each of the three perpendicular cavities. We selected these residues by visual inspection of the structures in PyMOL. We performed the cavity calculations on 20 structures extracted from each trajectory and each structural state identified from the clustering of the contact score. We selected the 20 structures with the lowest mean squared pairwise distance to other frames in the cluster. As a reference for comparing calculated tunnels, we included calculations on the starting models of the open and closed states.

We monitored the cavities with probe radii of 1.2 Å designed to resemble the size of a water molecule. As CAVER parameters, we used a shell radius of 13 Å, a shell depth of 4 Å, and a clustering threshold of 12 Å. We clustered the cavities using the hierarchical average-link clustering method of CAVER by which two cavities are clustered based on the dissimilarity of their pathways in the protein, calculated by dividing each tunnel in a sequence of *N* consecutive points starting from the reference site and then calculating their pairwise distance. We used in-house Python and R scripts to i) follow the evolution of the cavities during the MD trajectories, calculate the average radius and length of tunnels, and to identify the residues lining the cavities. We used PyMOL to plot the cavities and graphically visualize them. Here, we focused on comparing the top ten clusters of tunnels identified by CAVER, ranked based on the quantification of their tunnel throughputs.

### Analysis of membrane properties and protein-lipid contacts

We used *LipidDyn*^28^ to calculate the lipid bilayer properties and dynamics and protein-lipid contacts. During the simulation, we monitored the lipid transbilayer (i.e., scrambling) movements. The implementation of the lipid transbilayer module was officially included in the *LipidDyn* GitHub repository on 14/07/2023 (https://github.com/ELAB/LipidDyn/pull/157). *LipidDyn* uses the *LeafletFinder* class of MDAnalysis^61^ to identify the leaflet of the bilayer and the lipids belonging to them, considering atoms in their headgroups as representative atoms of each molecule. *LipiDdyn* employs a reimplementation of *FATSLiM* software to estimate the bilayer thickness as the distance vector between neighborhood-averaged coordinates of each lipid and its neighbors in the opposite leaflet, using a cut-off distance of 60 Å. Furthermore, the calculation of APL is performed by a neighbor search of each lipid and computation of a Voronoi tessellation. As a note, we did not discuss in in this study the analyses of membrane curvature since we could not estimate relevant differences among ATG9A and ATG9A_N99glyco_ and larger bilayers would be required. *LipidDyn* incorporates an analysis module to track and analyze the lipid transbilayer movements. This tool uses the positions of the leaflet groups calculated from the *LeafletFinder* class to estimate the shape of the surface of the two leaflets to provide a reference for calculating the lipid transbilayer movements. The surface estimation is performed through a linear regression model, using four parameters as the polynomial degree of the equation. The tool then tracks the positions of lipid headgroups, enabling the identification of lipids that undergo trans-bilayer movements by using distance cutoffs from the two leaflets. We considered lipids that undergo even partial trans-bilayer movements using as a distance cut off the 20% of the distance between the two leaflets of the bilayer. Furthermore, it calculates the contacts of the identified lipids with proteins using a distance cutoff of 6 Å.

### Cell lines and Culture

Immortalized human embryonic kidney cells (HEK293) cell line were obtained from the American Type Culture Collection (ATCC, ref. CRL-1573). ATG9A KO HEK293 cell line was obtained from Richard Youle’s lab. Cells were grown in Dulbecco’s modified eagle’s medium (DMEM) high glucose (Gibco #31966-021) supplemented with 10% fetal bovine serum (Gibco #10270-106). Cell lines were grown at 37°C in a humidified incubator containing 5% CO_2_.

### Plasmid and DNA transfection

V5-ATG9A plasmid was generated by insertion of human ATG9A sequence into a V5-TurboID-NES pCDNA3 plasmid (Addgene #107169) C-terminal to the V5-tag using the NEBuilder HiFi DNA Assembly Cloning Kit (New England Biolabs (NEB), #E5520S) according to the user guidelines. Subsequently, the sequence encdoding TurboID-NES was deleted by mutagenesis using the Platinum SuperFi II DNA Polymerase system (Thermo Scientific #12361010). ATG9A mutants (N99D and N99A) were generated by site-directed mutagenesis using the Platinum SuperFi II DNA Polymerase system and confirmed by sequencing (Eurofins). The cells were transfected with DNA using polyethylenimine (PEI) (PolySciences, #23966-100) in OptiMEM (ThermoFisher) according to the manufacturer guidelines. Oligonucleotides used in this study were obtained from TAG Copenhagen and their sequences are listed in TableS3. The DNA sequence of V5-ATG9A is listed in TableS4.

### Antibodies

Rabbit ATG9A (Cell Signaling Technology, #13509), rabbit LC3B (Cell Signaling Technology, #2775), mouse beta Actin (C4) (Santa Cruz Biotechnology, #sc-47778), rabbit SQSTM1/p62 (MBL, #PM045).

### Starvation assay

Cells were seeded in 6 cm plates at 3×10^5^ cells/mL. 24 hours post transfection the cells were washed twice in Earle’s Balanced Salt Solution (EBSS; Sigma-Aldrich, E2888) medium and then incubated in EBSS medium for 4 h with or without 200 nM Bafilomycin A1. Cells were washed three times in PBS before being transferred to 350 µL RIPA lysis buffer (50mM Tris-HCl pH 7.4, 150mM NaCl, 0.1% SDS, 0.5% sodium deoxycholate, 1% Triton X-100, 5mM EDTA) containing protease inhibitor cocktail, NaV, NaF, and β-glycerophosphate, using a cell scraper. Lysis was performed by incubating cells on ice for 30 min. The lysate was centrifuged at 15.000xG for 15 min at 4 °C and the supernatant was transferred to a new tube. The protein concentration was determined using the DC protein assay (BioRad).

### Western blot

Samples for western blots were mixed with NuPAGE SDS-loading buffer and NuPAGE reducing agent. Samples were incubated at 65 °C for 15 min prior to separation by electrophoresis using a 15% SDS polyacrylamide gel and subsequently electro transferred to a nitrocellulose membrane (BioRad). The membrane was blocked by incubation in PBS-T (DPBS, 0.1 % Tween-20) containing 5% (w/v) nonfat dry milk powder for 1 h. The membrane was incubated in primary antibody (1:1000 dilution) for 1 h at room temperature or overnight at 4 °C. The membrane was washed 5 times for 5 min with PBS-T to removed unbound primary antibody and incubated for 1 h at room temperature with the corresponding horseradish peroxidase-conjugated secondary antibody (Goat Anti-rabbit, #1706515 or Anti-mouse, 1706516, IgG (H+L)-HRP Conjugate Bio-Rad) at a 1:3000 dilution. The membrane was subsequently washed 5 times for 5 min to remove unbound antibodies in PBS-T. The membranes were developed using chemiluminescence (Amersham™ ECL Select™ Western Blotting Detection Reagent, Cytiva) and images were acquired using a ChemiDoc™ Imaging system, Bio-Rad.

To normalize for protein loading, the membrane has been incubated with a mouse monoclonal anti β-actin antibody (Santa Cruz Biotechnology, #sc-47778). Relative band intensities were determined using the ImageLab software (Bio-Rad). To account for variations in basal LC3-II and SQSTM1/p62 levels under steady-state conditions, autophagy flux was quantified using specific normalization ratios. The synthesis ratio was calculated by comparing the levels of selected autophagy markers in the presence of both an autophagy inducer and a degradation inhibitor to their levels with the inhibitor alone, representing the rate of autophagosome formation. The degradation ratio was determined by comparing marker levels when both the inducer and inhibitor were applied to their levels with the inducer alone, providing a measure of autophagosome turnover.

## Conclusions

This study used microsecond all-atom molecular dynamics simulations to investigate the effects of N99 glycosylation on ATG9A, offering insights into its conformational dynamics and lipid scrambling mechanism. The ATG9A hydrophilic central cavity supports phospholipid headgroup insertion, reorientation, and partial transbilayer movements, consistent with lipid scrambling activity observed in experimental assays. Of note, the MD simulations discussed here included the fully N99-glycosylated form of ATG9A with a complex-type *N-*glycan, a common modification in human proteins. However, it is important to note that in the cells, ATG9A trimers might exist as a mixture of monomers with varying levels of glycosylation. This potential heterogeneity should be considered when interpreting experimental results, as it may influence the structural and functional properties of the trimers. Since the loop containing N99 and the *N-*glycans are unresolved in the ATG9A cryo-EM structures and their heterogeneity complicates experimental characterization, our simulations provide valuable insights into the structural and functional consequences of N-glycosylation on ATG9A. Intrinsic flexibility is a conserved feature of ATG9 proteins, as both ATG9A and ATG9B exhibit significant conformational plasticity, suggesting this is an inherent characteristic crucial for their function. A key determinant of this flexibility is the HINGE region (i.e., TMH1, the reentrant membrane helix 1, TMH3, and TMH4), whose structural differences between cryo-EM structures and AlphaFold models further emphasize its role in modulating ATG9A dynamics and lipid interactions^62^. Our findings support the idea that this region directly impacts lipid handling and transport, aligning with its proposed role in autophagy. Glycosylation may fine-tune this inherent flexibility, rather than being strictly required for lipid scramblase activity. While Chiduza et al. demonstrated that lipid scrambling occurs in the absence of glycosylation^62^, we show that glycosylation might modulate ATG9A dynamics. Our results further support this notion, revealing that *N*-glycosylation strengthens cooperative interactions between protomer conformations, facilitating lipid insertion and partial traversal within the central cavity. This aligns with the proposed mechanism of ATG9A role in autophagy, where it aids lipid redistribution across the phagophore membrane to support elongation and autophagosome biogenesis. Our findings, however, suggest that abolishing N-glycosylation through mutagenesis in an experimental assay for autophagy flux were not sufficient to identify a marked change in the protein functionality, suggesting that future *in-vitro* experiments based on lipid scramblase assays could better clarify the effects of the glycosylation. Furthermore, varying levels of ATG9A functionality could impact autophagosome size, highlighting the need for specific experiments to explore this relationship. On the other hand, we observed partial phospholipid traversal at protomer interfaces, suggesting that complete transbilayer movements may require enhanced sampling techniques or larger membrane models to incorporate effects of curvature and biophysical membrane properties. Finally, our simulations show that ATG9A can adopt asymmetric conformations at protomer interfaces, in contrast with symmetric configurations observed in cryo-EM structures. These findings could guide further analysis of cryo-EM datasets to uncover structural heterogeneity. Our work emphasizes the importance of incorporating glycosylation in computational studies of ATG9A dynamics and lipid scrambling. The methods developed here can be extended to study other scramblases and flippases, advancing our understanding of lipid transport mechanisms in various cellular contexts.

## Author Contributions (CRediT Classification)

Conceptualization: ML, EP. Data Curation: ML, MU. Formal Analysis: ML, MU, HBM, SEE Funding Acquisition: EP, MJ. Investigation: ML, MU, EP. Methodology: ML, MU, HBM, CBB, EF,KM, NT, MJ. Project administration: EP. Resources: EP, MJ. Supervision: EP, ML. Validation: ML, MU. Visualization: ML, MU. Writing – Original Draft: ML, MU, EP. Writing – Review and Editing: All the coauthors.

## Declaration of generative AI and AI-assisted technologies in the writing process

During the preparation of this work the author(s) used OpenAI ChatGPT 3.5 to improve the language of the manuscript. After using this service, the authors reviewed and edited the content as needed and take full responsibility for the content of the published article.

### Acknowledgments

Our research has been supported by Danmarks Grundforskningsfond (DNRF125) to MJ and EP, along with by Danmarks Frie Forskningsfond, Natural Science, Project 1 (102517) and NovoNordisk Fonden Bioscience and Basic Biomedicine (NNF20OC0065262) to EP. Part of the calculations described in this paper were performed thanks to DECI-PRACE 17th to ML for calculations on Archer2 (UK) and to the Danish HPC Infrastructure Computerome2 access. Part of the calculations have been supported by a EuroHPC Benchmark Access Grant (EHPC-BEN-2023B02-010) and a EuroHPC Regular Grant (EHPC-REG-2023R01-051) on Discoverer. The Novo Nordisk Foundation Center for Protein Research is supported financially by the Novo Nordisk Foundation (NNF14CC0001). NMIT acknowledges support from an NNF Hallas-Møller Ascending Investigator grant (NNF23OC0081528). NMIT is also a member of the Integrative Structural Biology Cluster (ISBUC) at the University of Copenhagen. We would like to thank Isha Raj for the helpful scientific discussions in the early stages of the project.

