## Supplementary Figures for "Role of *N*-glycosylation as a determinant of ATG9A conformations and activity"

RMSD Cα atoms of the swapped helices of each protomer of ATG9A

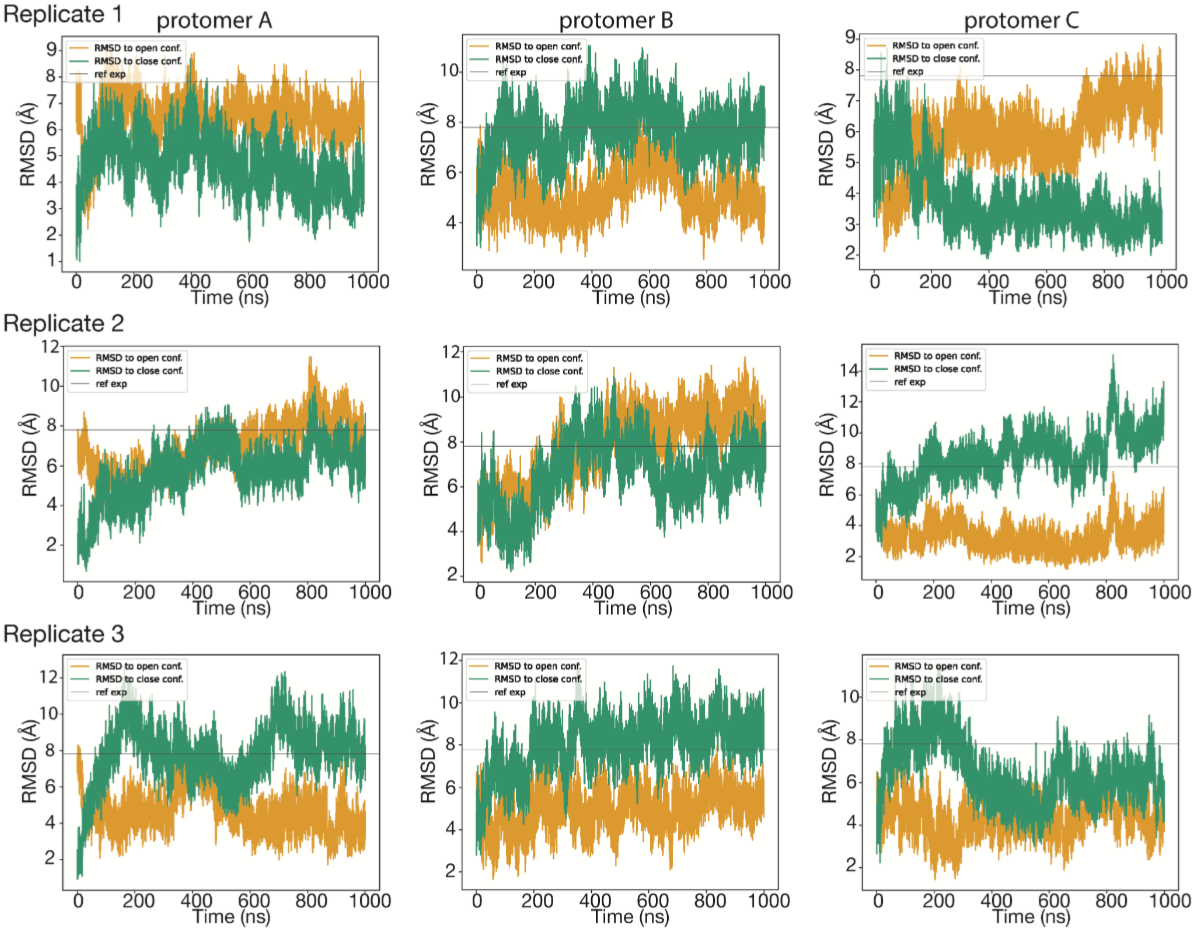


RMSD Cα atoms of the swapped helices of each protomer of $ATG9A_{N99glyco}$

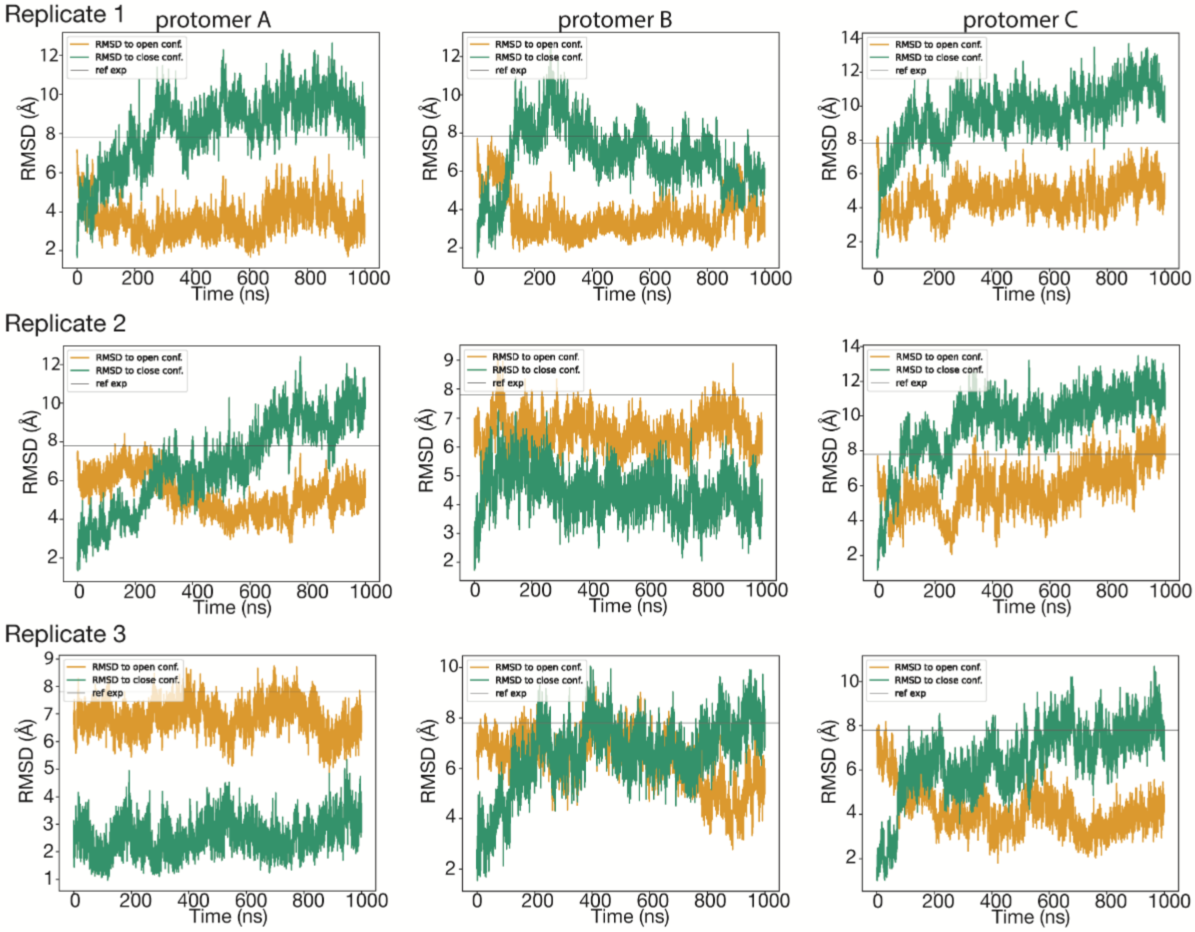

**Figure S1. ATG9A and ATG9A$_{N99glyco}$ starting from the closed state transition towards the open state of at least one protomer**. Line plots of the RMSD values of the Cα atoms of the domain-swapped transmembrane helices of each protomer (protomer A, B and C) relative to the experimental cryo-EM structures of the open (yellow, PDB ID 6WQZ, PMID: 32610138) and closed (green, PDB ID 7JLP, PMID: 33106659) state of ATG9A. The RMSD has been calculated for the replicate 1-3 (left, middle and right panels, respectively) of ATG9A (upper panels) and ATG9A$_{N99glyco}$ (upper panels). For reference, the RMSD between the closed and open state cryo-EM structures is approximately 7.8 Å and indicated in the plots by a gray line.

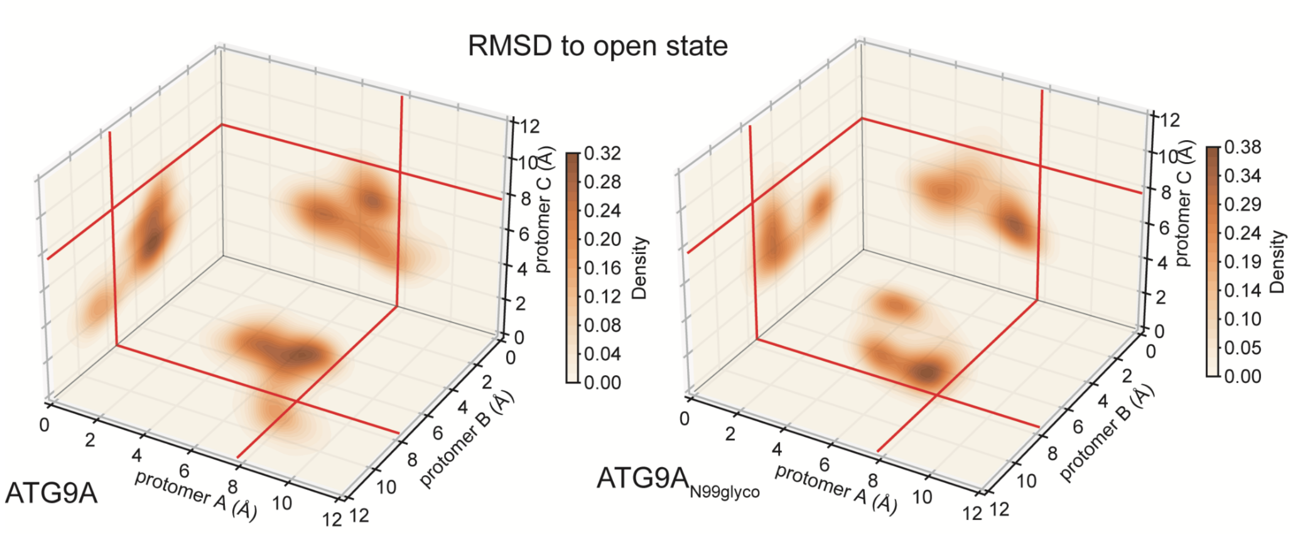


**Figure S2 ATG9A and ATG9A$_{N99glyco}$ assumes asymmetric open states of the protomers**. Density plots of the RMSD values calculated for ATG9A and ATG9A$_{N99glyco}$ on the Cα atoms of the domain-swapped transmembrane helices (TMH3 and TMH4) of each protomer (named protomer A, B and C) relative to the experimental cryo-EM structures of the open state of ATG9A (PDB ID 6WQZ, PMID: 32610138). The plots show the projection of the density for the different protomers along the concatenated trajectories of the replicates. For reference, the RMSD between the closed and open state is approximately 7.8 Å and indicated in the plots by a red line.

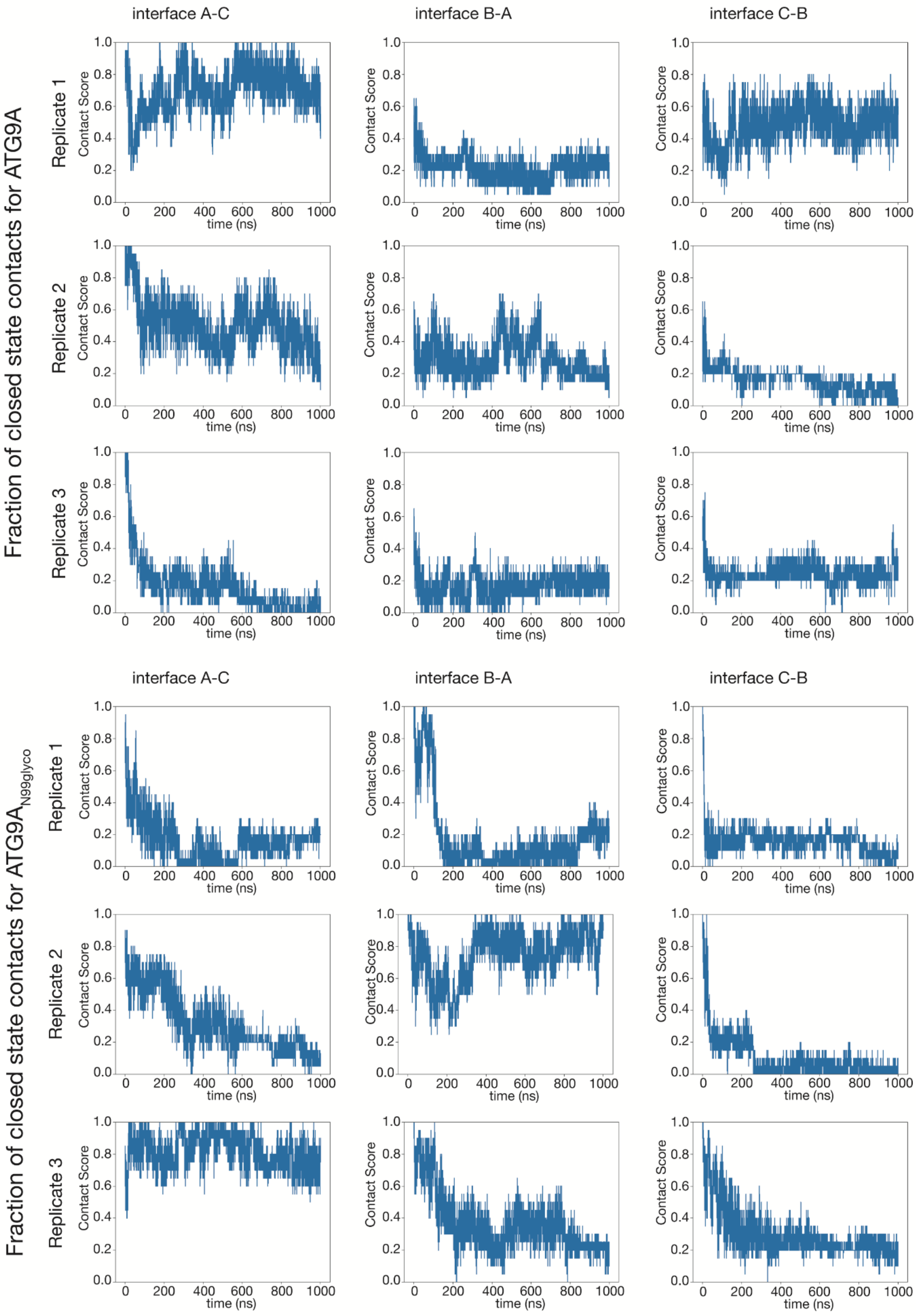
Fraction of closed state contacts for ATG9A
Fraction of closed state contacts for $ATG9A_{N99glyco}$
interface A-C
interface B-A
interface C-B
Replicate 1
Replicate 2
Replicate 3
Contact Score
time (ns)
0.0
0.2
0.4
0.6
0.8
1.0
0
200
400
600
800
1000

**Figure S3 ATG9A and ATG9A$_{N99glyco}$ undergo asymmetric closed-open conformational changes.** Line plots of the contact score values calculated for ATG9A and ATG9A$_{N99glyco}$ by monitoring the fraction of protein-protein contacts among the domain-swapped TMH3–4 helices of each protomer and other regions of ATG9A that are present only in the cryo-EM structure of the closed state of ATG9A (PDB ID 7JLP, PMID: 33106659). The collective contact score has been calculated for each protomer-protomer interface (interfaces AC, BA, and CB) for the replicate 1-3 (upper, middle and lower panels, respectively) of ATG9A (upper panels) and ATG9A$_{N99glyco}$ (upper panels).

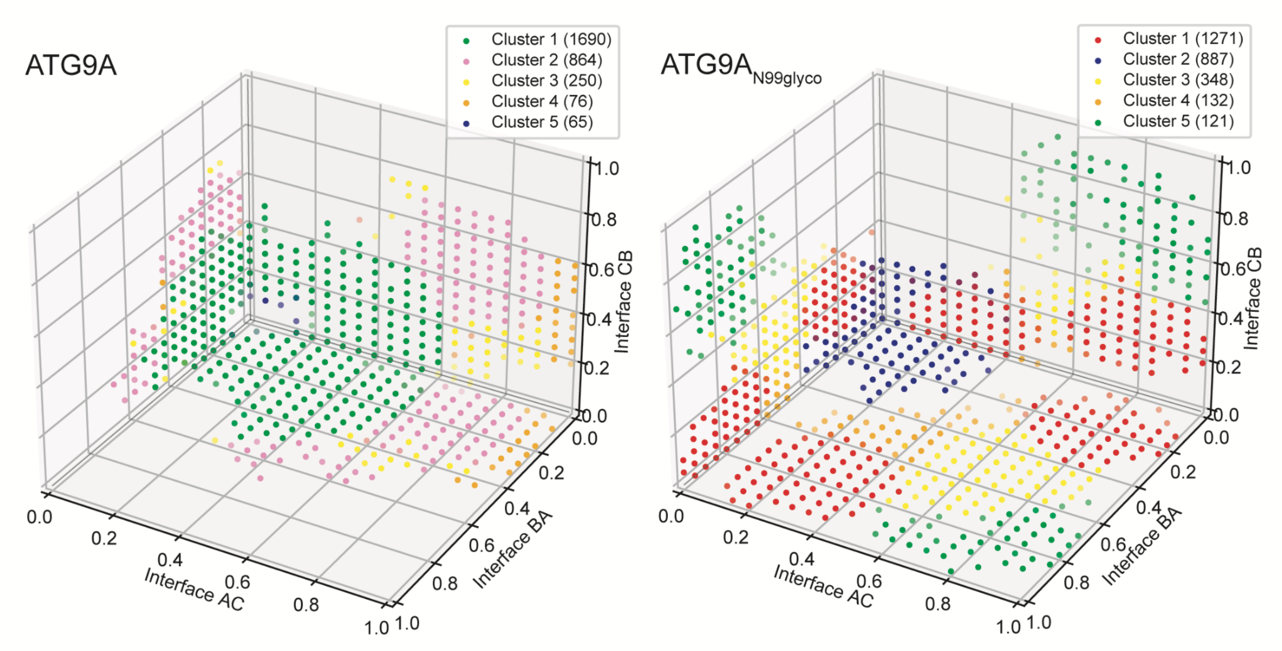


**Figure S4 N-glycosylation at N99 favors the extent of open conformations in ATG9A**. The three-dimensional plots show the results of the cluster analysis calculated on the concatenated trajectories of ATG9A and ATG9A$_{N99glyco}$ using Euclidean distance metric based on the contact scores of each protomer-protomer interface AC, BA, and CB. A quality-threshold algorithm was employed with a threshold of 0.45, discarding clusters with less than 60 frames (less 2% of the total). Each dot represents a structure from the concatenated trajectories colored according to the cluster ID. Cluster 1 of ATG9A included partially open states, including around 56% of the concatenated trajectory frames. ATG9A$_{N99glyco}$ exhibits a higher occurrence than ATG9A of states with open conformations at all protomer interfaces, as represented by cluster 2, encompassing around 30% of the concatenated trajectory frames.

### Close conformation contact score

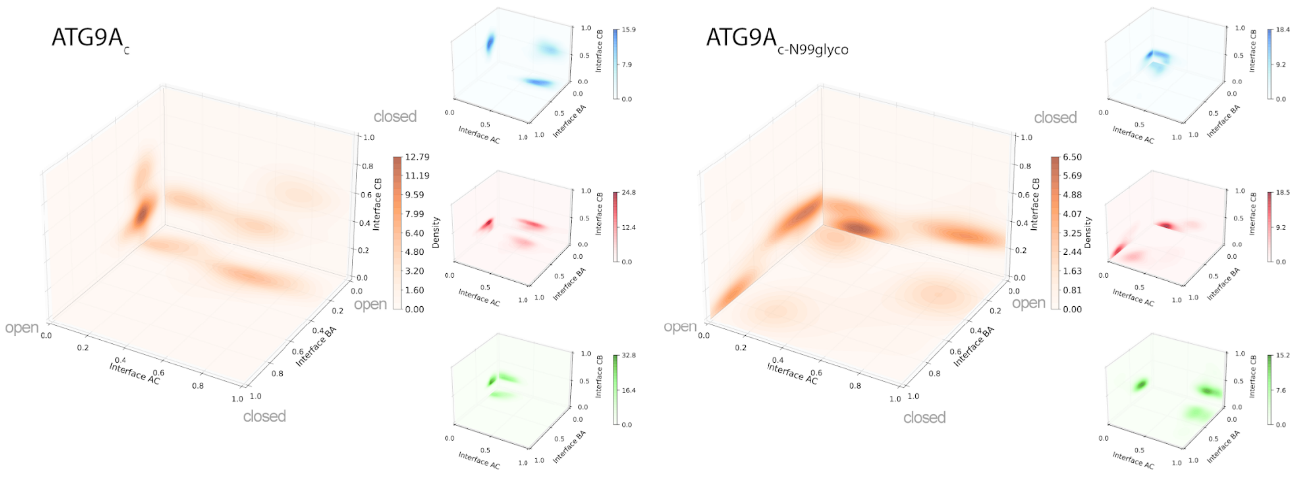


### RMSD to close referece structure

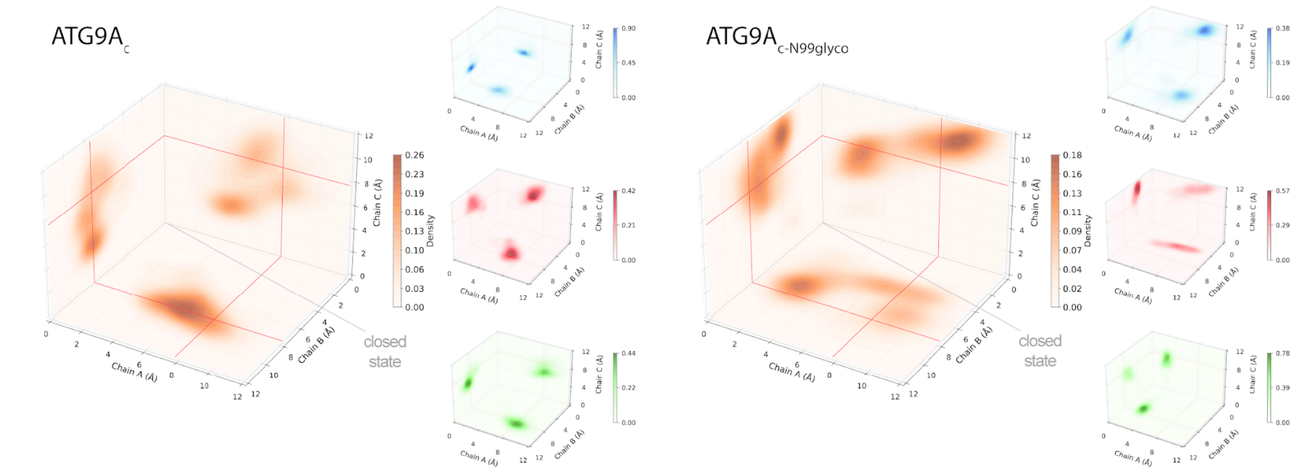


### RMSD to open reference structure

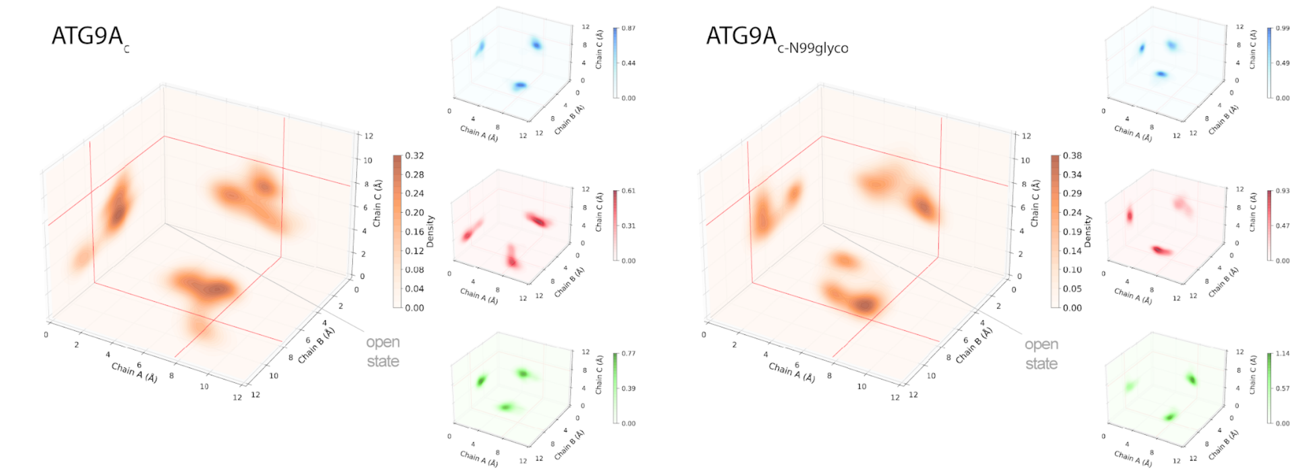

**Figure S5 Analysis of behavior of the single MD replicates of ATG9A and ATG9A$_{\mathbf{N99glyco}}$.** A) Cluster analysis using the collective contact score values calculated by monitoring the occurrence of protein-protein contacts within the regions RMH1-turn-helix α9 and TMH3–4 that are present only in the experimental cryo-EM structure of the closed state of ATG9A. A quality-threshold algorithm was used with a threshold of 0.45, discarding clusters with less than 60 frames (less 2% of the total) (upper panels). B-C) RMSD of domain-swapped transmembrane helices relative to the experimental cryo-EM structure of the closed (middle panels) and open (lower panels) state.

### Replicate1

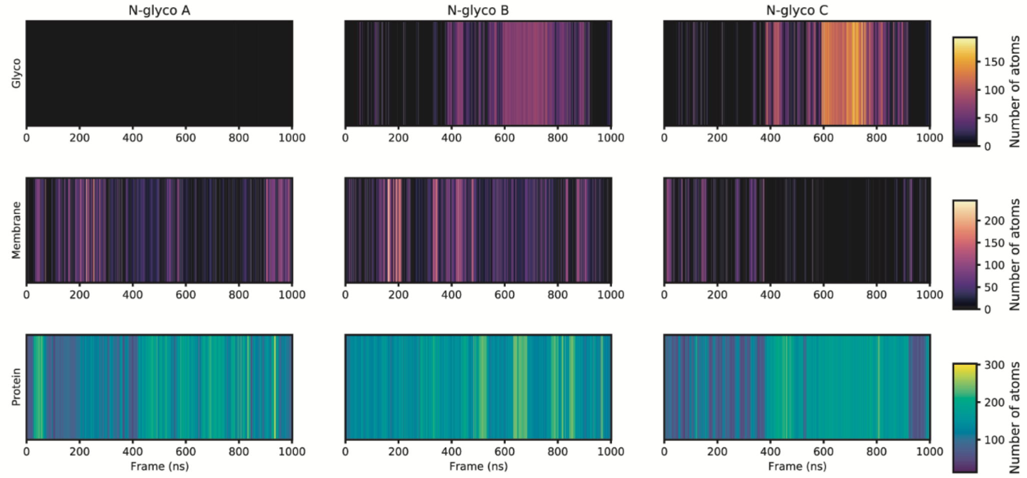


### Replicate2

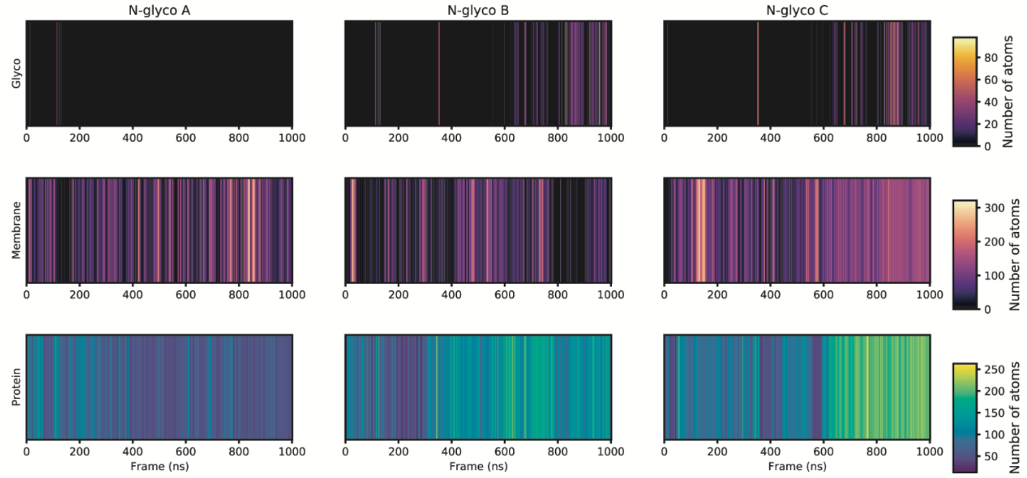


### Replicate3

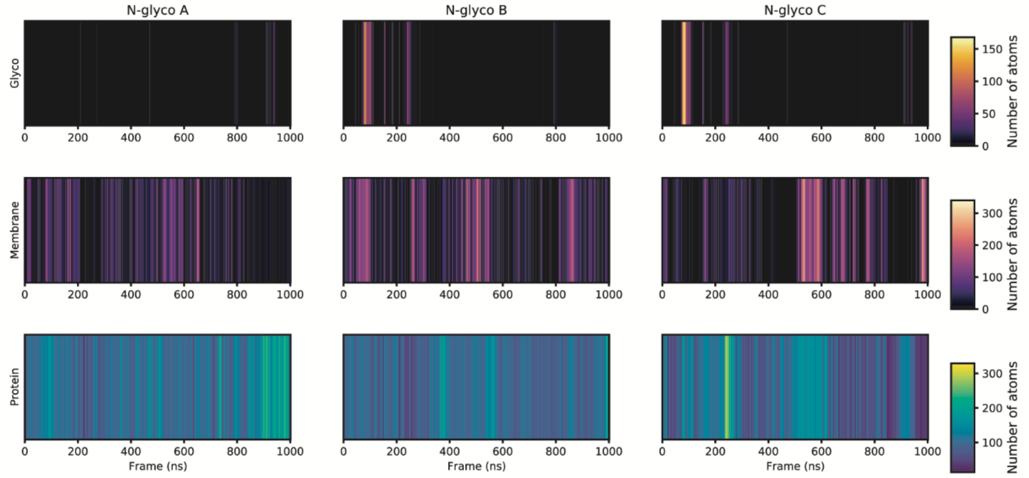

**Figure S6 Glycan, protein, and lipid atoms in the surroundings of the three N-glycosylations form heterogeneous patterns of transient interactions.** The heatmaps show the number of glycan, protein, and lipid atoms in the surroundings (6 Å distance cutoff) of the three N-glycosylations for the three replicates of ATG9A$_{N99glyco}$.

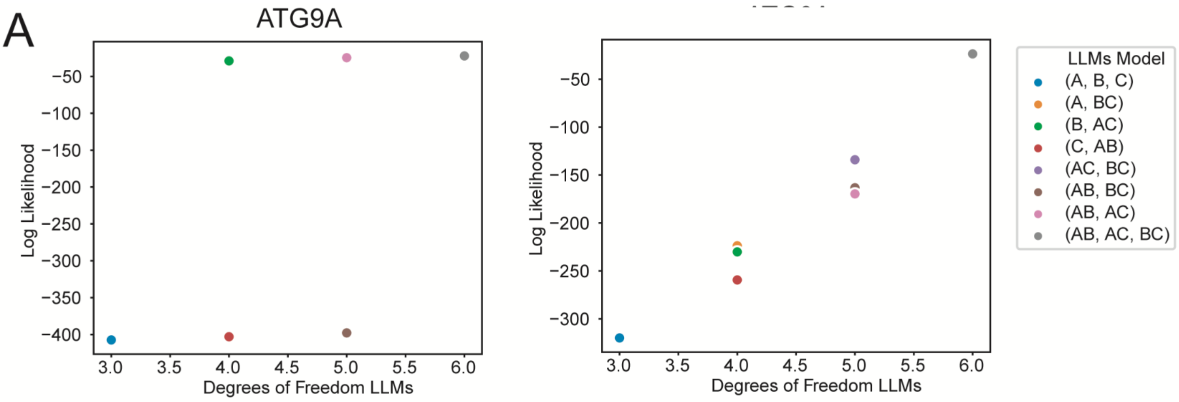


B

ATG9A

| Degree of freedom LLMs | Degree of freedom LLMs | meandiff. | p-value adj. | Tukey's HSD |
|---|---|---|---|---|
| 3 | 4 | -258.4916 | 0.9 | False |
| 3 | 5 | -516.9831 | 0.7296 | False |
| 3 | 6 | -769.8727 | 0.6247 | False |
| 4 | 5 | -258.4916 | 0.8718 | False |
| 4 | 6 | -511.3811 | 0.7348 | False |
| 5 | 6 | -252.8896 | 0.9 | False |

ATG9A$_{N99glyco}$

| Degree of freedom LLMs | Degree of freedom LLMs | meandiff | p-value adj. | Tukey's HSD |
|---|---|---|---|---|
| 3 | 4 | -164.2396 | 0.0651 | False |
| 3 | 5 | -328.4792 | 0.0059 | True |
| 3 | 6 | -592.5287 | 0.0013 | True |
| 4 | 5 | -164.2396 | 0.0207 | True |
| 4 | 6 | -428.2891 | 0.0022 | True |
| 5 | 6 | -264.0495 | 0.0131 | True |

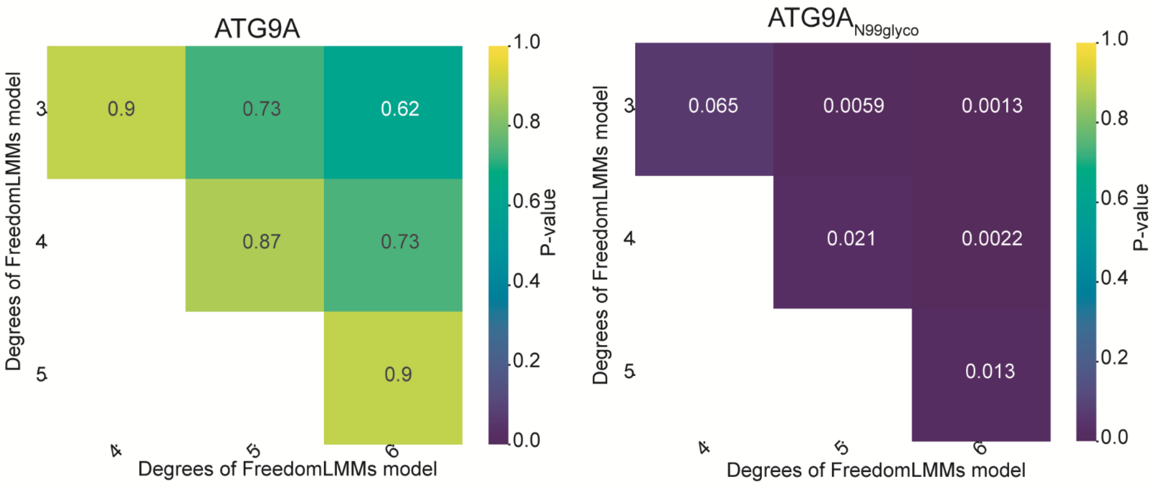

**Figure S7 N-glycosylation promotes cooperativity in the closed-open conformational changes of ATG9A.** (A) Log-Linear Models (LLMs) analysis illustrating the relationships among open/closed states of the protomers (A, B, C) of ATG9A (left panels) and $ATG9A_{N99glyco}$ (right panels). The upper panels show the degrees of freedom for the different LLMs and their log-likelihoodness in fitting the contact score data, highlighting differences in the fit of independence models between ATG9A and $ATG9A_{N99glyco}$. We tested complete independence ([A, B, C]), joint dependence ([A,BC], [B,AC], [C,AB]), conditional independence ([AC,BC], [AB,BC], [AB,AC]) and homogeneous association model (i.e., [AB, AC, BC]). B) Tukey Honestly Significant Difference (HSD) tests to analyze the deviance and log-likelihood data, providing insights into the statistical significance of the model comparisons. The lower panels show as heatmaps the p-value calculated to assess the significance of the comparison between LLMs models with different degrees of freedom.

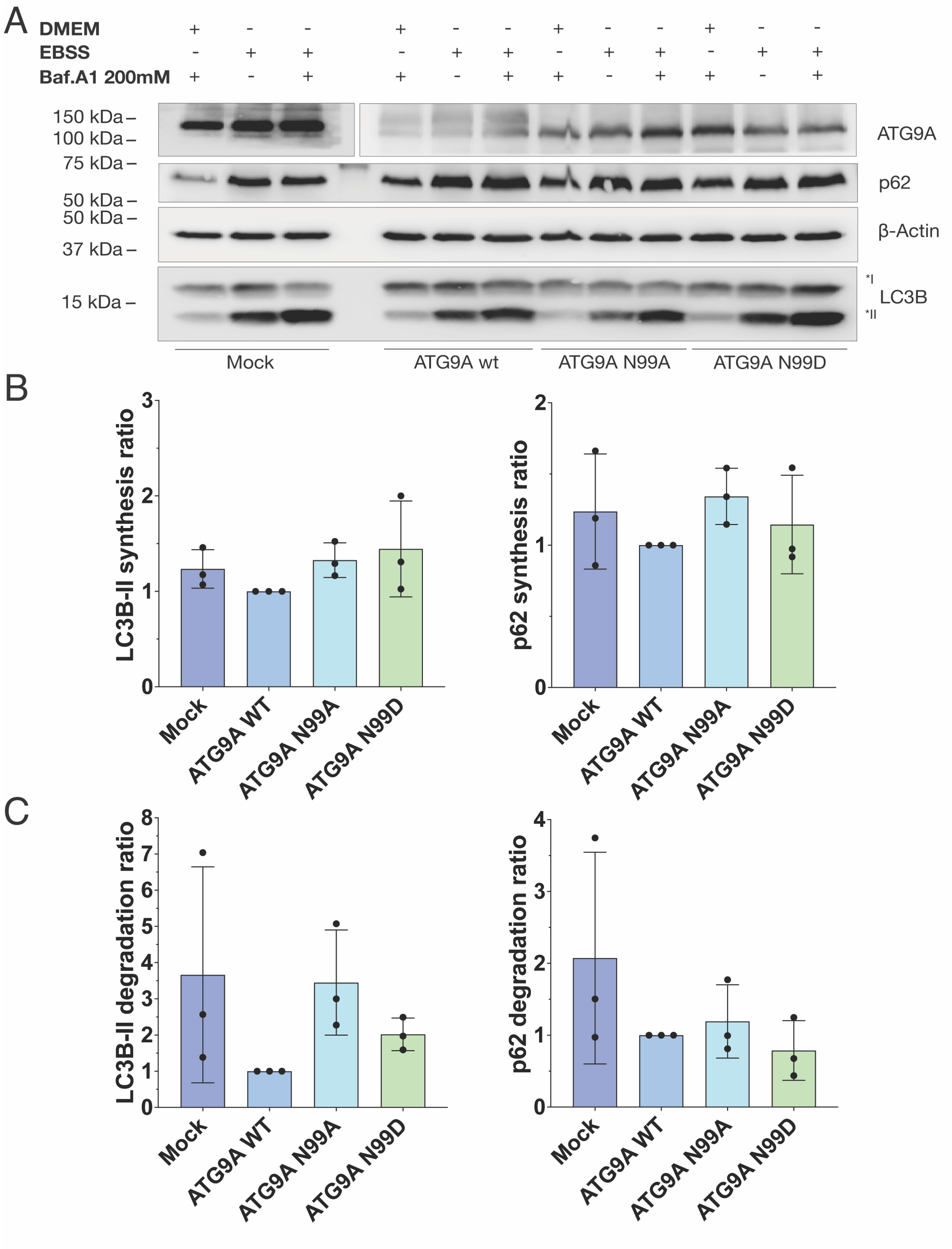


**Figure S8. Autophagy flux analysis on the control HEK293 cell line.** (A) Western blot analysis of the LC3-II and p62 levels of ATG9A WT, ATG9A$^{N99A}$ and ATG9A$^{N99D}$ under starvation (EBSS medium) or rich-nutrient conditions (DMEM medium), in the presence and absence of bafilomycin A1, a lysosomal degradation inhibitor. $ATG9A^{N99A}$ and $ATG9A^{N99D}$ samples were both loaded on a 15% gel along with ATG9A WT and the Mock condition (i.e., no overexpression). LC3-II and SQSTM1/p62 levels have been normalized with the housekeeping protein β-Tubulin levels in the same sample. (B) Analysis of the autophagosome formation rate (synthesis rate) following LC3-II (left) or SQSTM1/p62 (right) levels. The synthesis rate has been calculated as the rate of LC3-II (or SQSTM1/p62) levels in the presence of the inducer (i.e., EBSS) and the inhibitor (i.e., Bafilomycin A1) divided by the LC3-II (or SQSTM1/p62) level in the presence of the inhibitor alone. (C) Analysis of the autophagosome degradation rate following LC3-II (left) or SQSTM1/p62 (right) levels. The degradation rate has been calculated as the rate of LC3-II (or SQSTM1/p62) levels in the presence of the inducer (i.e., EBSS) and the inhibitor (i.e., Bafilomycin A1) divided by the LC3-II (or SQSTM1/p62) level in the presence of the inducer alone.
