## Supplementary material for "Role of *N*-glycosylation as a determinant of ATG9A conformations and activity": Table S1

**Table S1. Similarity between the ensembles of contact score values of the interfaces AC/BA/CB defined by each cluster in ATG9A and ATG9A_N99glyco_.** As similarity measurements, we calculated the Jaccard similarity index and Weighted Jaccard similarity index to take into account the frequencies of the values. The results are in both cases a value between 0 and 1, where 1 indicates identical sets of values, and 0 indicates no similarity.

| ***Cluster ID ATG9A (***% of the concatenated trajectory frames) | ***Cluster ID***  ***ATG9A_N99Glyco_ (***% of the concatenated trajectory frames) | ***Jaccard Similarity Index*** | ***Weighted Jaccard Similarity index*** |
| --- | --- | --- | --- |
| *1 (56%)* | *1 (42%)* | *0.006802721088* | *0.004750593824* |
|  | *2 (30%)* | *0.1549019608* | *0.08413967186* |
|  | *3 (12%)* | *0* | *0* |
|  | *4 (4%)* | *0.04246284501* | *0.01334816463* |
|  | *5 (4%)* | *0* | *0* |
|  | *6 (3%)* | *0* | *0* |
|  | *7 (2%)* | *0* | *0* |
| *2 (29%)* | *1 (42%)* | *0.03827751196* | *0.01232811759* |
|  | *2 (30%)* | *0* | *0* |
|  | *3 (12%)* | *0.02615694165* | *0.01084236864* |
|  | *4 (4%)* | *0.01259445844* | *0.007077856421* |
|  | *5 (4%)* | *0* | *0* |
|  | *6 (3%)* | *0.002747252747* | *0.00103626943* |
|  | *7 (2%)* | *0* | *0* |
| *3 (8%)* | *1 (42%)* | *0.1081730769* | *0.05259515571* |
|  | *2 (30%)* | *0* | *0* |
|  | *3 (12%)* | *0.07023411371* | *0.05467372134* |
|  | *4 (4%)* | *0.009523809524* | *0.005263157895* |
|  | *5 (4%)* | *0* | *0* |
|  | *6 (3%)* | *0.005747126437* | *0.002849002849* |
|  | *7 (2%)* | *0* | *0* |
| *4 (3%)* | *1 (42%)* | *0.04266666667* | *0.01737160121* |
|  | *2 (30%)* | *0* | *0* |
|  | *3 (12%)* | *0* | *0* |
|  | *4 (4%)* | *0* | *0* |
|  | *5 (4%)* | *0* | *0* |
|  | *6 (3%)* | *0.02941176471* | *0.01714285714* |
|  | *7 (2%)* | *0* | *0* |
| *5 (2%)* | *1 (42%)* | *0* | *0* |
|  | *2 (30%)* | *0.1052631579* | *0.04730473047* |
|  | *3 (12%)* | *0* | *0* |
|  | *4 (4%)* | *0* | *0* |
|  | *5 (4%)* | *0* | *0* |
|  | *6 (3%)* | *0* | *0* |
|  | *7 (2%)* | *0* | *0* |
