## Supplementary material for "Role of *N*-glycosylation as a determinant of ATG9A conformations and activity": Table S2

**Table S2.**  **Analysis using LipidDyn of the CHL1 lipids showing partial trans-bilayer movements for each replica of ATG9A and ATG9A_N99glyco_.**

| **ATG9A** | | | **ATG9A_N99glyco_** | | |
| --- | --- | --- | --- | --- | --- |
| **Replicate 1** | **Replicate 2** | **Replicate 3** | **Replicate 1** | **Replicate 2** | **Replicate 3** |
| 523, 524, 525, 526, 527, 528, 529, 530, 531, 532, 533, 534, 535, 536, 537, 538, 539, 540, 541, 542, 543, 544, 545, 546, 547, 548, 549, 550, 551, 552, 553, 554, 555, 556, 557, 558, 559, 560, 561, 562, 566, 567, 568, 569, 570, 571, 572, 573, 574, 575, 576, 577, 578, 579, 581, 582, 583, 584, 585, 586, 587, 588, 589, 590, 591, 592, 593, 594, 595, 596, 597, 598, 599, 600, 601, 602, 603, 604, 605, 606, 610, 611, 612, 613, 614, 615, 616, 617, 618, 619, 620, 621, 622, 623, 624, 625, 626, 627, 628, 629, 630, 631, 632, 633, 634, 635, 636, 637, 638, 639, 640, 641, 642, 643, 644, 645, 646, 647, 648, 649, 653, 654, 655, 656, 657, 658, 659, 660, 661, 662, 663, 664, 665, 666, 667, 668, 669, 670, 671, 672, 673, 674, 675, 676, 677, 678, 679, 680, 681, 682, 683, 684, 685, 686, 687, 688, 689, 690, 691, 692, 696, 697, 698, 699, 700, 701 | 523, 524, 525, 526, 527, 528, 529, 530, 531, 532, 533, 534, 535, 536, 537, 538, 539, 540, 541, 542, 543, 544, 545, 546, 547, 548, 549, 550, 551, 552, 553, 554, 555, 556, 557, 558, 559, 560, 561, 562, 563, 564, 565, 566, 567, 568, 569, 570, 571, 572, 573, 574, 575, 576, 577, 578, 579, 580, 581, 582, 583, 584, 585, 586, 587, 588, 589, 590, 591, 592, 593, 594, 595, 596, 597, 598, 599, 600, 601, 602, 603, 604, 605, 606, 607, 608, 609, 610, 611, 612, 613, 614, 615, 616, 617, 618, 619, 620, 621, 622, 623, 624, 625, 626, 627, 628, 629, 630, 631, 632, 633, 634, 635, 636, 637, 638, 639, 640, 641, 642, 643, 644, 645, 646, 647, 648, 649, 650, 651, 652, 653, 654, 655, 656, 657, 658, 659, 660, 661, 662, 663, 664, 665, 666, 667, 668, 669, 670, 671, 672, 673, 674, 675, 676, 677, 678, 679, 680, 681, 682, 683, 684, 685, 686, 687, 688, 689, 690, 691, 692, 693, 694, 695, 696, 697, 698, 699, 700, 701 | 523, 524, 525, 526, 527, 528, 529, 530, 531, 532, 533, 534, 535, 536, 537, 538, 539, 540, 541, 542, 543, 544, 545, 546, 547, 548, 549, 550, 551, 552, 553, 554, 555, 556, 557, 558, 559, 560, 561, 562, 563, 564, 565, 566, 567, 568, 569, 570, 571, 572, 573, 574, 575, 576, 577, 578, 579, 580, 581, 582, 583, 584, 585, 586, 587, 588, 589, 590, 591, 592, 593, 594, 595, 596, 597, 598, 599, 600, 601, 602, 603, 604, 605, 606, 607, 608, 609, 610, 611, 612, 613, 614, 615, 616, 617, 618, 619, 620, 621, 622, 623, 624, 625, 626, 627, 628, 629, 630, 631, 632, 633, 634, 635, 636, 637, 638, 639, 640, 641, 642, 643, 644, 645, 646, 647, 648, 649, 650, 651, 652, 653, 654, 655, 656, 657, 658, 659, 660, 661, 662, 663, 664, 665, 666, 667, 668, 669, 670, 671, 672, 673, 674, 675, 676, 677, 678, 679, 680, 681, 682, 683, 684, 685, 686, 687, 688, 689, 690, 691, 692, 693, 694, 695, 696, 697, 698, 699, 700, 701 | 523, 524, 525, 526, 527, 528, 529, 530, 531, 532, 533, 534, 535, 536, 537, 538, 539, 540, 541, 542, 543, 544, 545, 546, 547, 548, 549, 550, 551, 552, 553, 554, 555, 556, 557, 558, 559, 560, 561, 562, 563, 564, 565, 566, 567, 568, 569, 570, 571, 572, 573, 574, 575, 576, 577, 578, 579, 580, 581, 582, 583, 584, 585, 586, 587, 588, 589, 590, 591, 592, 593, 594, 595, 596, 597, 598, 599, 600, 601, 602, 603, 604, 605, 606, 607, 608, 609, 610, 611, 612, 613, 614, 615, 616, 617, 618, 619, 620, 621, 622, 623, 624, 625, 626, 627, 628, 629, 630, 631, 632, 633, 634, 635, 636, 637, 638, 639, 640, 641, 642, 643, 644, 645, 646, 647, 648, 649, 650, 651, 652, 653, 654, 655, 656, 657, 658, 659, 660, 661, 662, 663, 664, 665, 666, 667, 668, 669, 670, 671, 672, 673, 674, 675, 676, 677, 678, 679, 680, 681, 682, 683, 684, 685, 686, 687, 688, 689, 690, 691, 692, 693, 694, 695, 696, 697, 698, 699, 700, 701, 702 | 523, 524, 525, 526, 527, 528, 529, 530, 531, 532, 533, 534, 535, 536, 537, 539, 540, 541, 542, 543, 544, 545, 546, 547, 548, 549, 550, 551, 552, 553, 554, 555, 556, 557, 558, 559, 560, 561, 562, 563, 564, 565, 566, 567, 568, 569, 570, 571, 572, 573, 574, 575, 576, 577, 578, 579, 580, 581, 582, 583, 584, 585, 586, 587, 588, 589, 590, 591, 592, 593, 594, 595, 596, 597, 598, 599, 600, 601, 602, 603, 604, 605, 606, 607, 608, 609, 610, 611, 612, 613, 614, 615, 616, 617, 618, 619, 620, 621, 622, 623, 624, 625, 626, 627, 628, 629, 630, 631, 632, 633, 634, 635, 636, 637, 638, 639, 640, 641, 642, 643, 644, 645, 646, 647, 648, 649, 650, 651, 652, 653, 654, 655, 656, 657, 658, 659, 660, 661, 662, 663, 664, 665, 666, 667, 668, 669, 670, 671, 672, 673, 674, 675, 676, 677, 678, 679, 680, 681, 682, 683, 684, 685, 686, 687, 688, 689, 690, 691, 692, 693, 694, 695, 696, 697, 698, 699, 700, 701, 702 | 523, 524, 525, 526, 527, 528, 529, 530, 531, 532, 533, 534, 535, 536, 537, 539, 540, 541, 542, 543, 544, 545, 546, 547, 548, 549, 550, 551, 552, 553, 554, 555, 556, 557, 558, 559, 560, 561, 562, 563, 564, 565, 566, 567, 568, 569, 570, 571, 572, 573, 574, 575, 576, 577, 578, 579, 580, 581, 582, 583, 584, 585, 586, 587, 588, 589, 590, 591, 592, 593, 594, 595, 596, 597, 598, 599, 600, 601, 602, 603, 604, 605, 606, 607, 608, 609, 610, 611, 612, 613, 614, 615, 616, 617, 618, 619, 620, 621, 622, 623, 624, 625, 626, 627, 628, 629, 630, 631, 632, 633, 634, 635, 636, 637, 638, 639, 640, 641, 642, 643, 644, 645, 646, 647, 648, 649, 650, 651, 652, 653, 654, 655, 656, 657, 658, 659, 660, 661, 662, 663, 664, 665, 666, 667, 668, 669, 670, 671, 672, 673, 674, 675, 676, 677, 678, 679, 680, 681, 682, 683, 684, 685, 686, 687, 688, 689, 690, 691, 692, 693, 694, 695, 696, 697, 698, 699, 700, 701, 702 |
| **Total** 163 | **Total** 179 | **Total** 179 | **Tota**l 180 | **Total** 180 | **Total** 180 |
