## Supplementary material for "Role of *N*-glycosylation as a determinant of ATG9A conformations and activity": Table S3

**Table S3. Primers used in this study.** Oligonucleotides used in this study were obtained from TAG Copenhagen.

| **Oligonucleotides** | **Forward primer** | **Reverse primer** |
| --- | --- | --- |
| Forward and Reverse primers to generate V5-TurboID-NES pCDNA3 compatible ATG9A fragment, using NEBuilder assembly | cagcaccgctATGGCGCAGTTTGACACTG | tgtctttgctTACCTTGTGCACCTGAGG |
| Forward and Reverse primers to generate ATG9A compatible V5-TurboID-NES pCDNA3 fragment, using NEBuilder assembly | cacaaggtaAGCAAAGACAATACTGTGCCTCTGAAGC | actgcgccatAGCGGTGCTGTCCAGGCC |
| Forward and Reverse primers to delete TurboID-NES sequence in V5-TurboID-NES pCDNA3, using Platinum SuperFi II DNA Polymerase system | TACCTTGTGCACCTGAGGGG | TAATAGCTCGAGCATGCATCTAGAGG |
| Forward and Reverse to generate N99A ATG9A mutant, using Platinum SuperFi II DNA Polymerase system | AGATGGTGGCCCACAGTCTTCACC | GACTGTGGGCCACCATCTTGTTGG |
| Forward and Reverse to generate N99D ATG9A mutant, using Platinum SuperFi II DNA Polymerase system | AGATGGTGGACCACAGTCTTCAC | ACTGTGGTCCACCATCTTGTTGG |
