## Supplementary material for "Role of *N*-glycosylation as a determinant of ATG9A conformations and activity": Table S4

**Table S4. DNA sequence of theV5-ATG9A construct used in this study.**

| DNA sequence of V5-ATG9A | ATGGGCAAGCCCATCCCCAACCCCCTGCTGGGCCTGGACAGCACCGCTATGGCGCAGTTTGACACTGAATACCAGCGCCTAGAGGCCTCCTATAGTGATTCACCCCCAGGGGAGGAGGACCTGTTGGTGCACGTCGCCGAGGGGAGCAAGTCACCTTGGCACCATATTGAAAACCTTGACCTCTTCTTCTCTCGAGTTTATAATCTGCACCAGAAGAATGGCTTCACATGTATGCTCATCGGGGAGATCTTTGAGCTCATGCAGTTCCTCTTTGTGGTTGCCTTCACTACCTTCCTGGTCAGCTGCGTGGACTATGACATCCTATTTGCCAACAAGATGGTGAACCACAGTCTTCACCCTACTGAACCCGTCAAGGTCACTCTGCCAGACGCCTTTTTGCCTGCTCAAGTCTGTAGTGCCAGGATTCAGGAAAATGGCTCCCTTATCACCATCCTGGTCATTGCTGGTGTCTTCTGGATCCACCGGCTTATCAAGTTCATCTATAACATTTGCTGCTACTGGGAGATCCACTCCTTCTACCTGCACGCTCTGCGCATCCCTATGTCTGCCCTTCCGTATTGCACGTGGCAAGAAGTGCAGGCCCGGATCGTGCAGACGCAGAAGGAGCACCAGATCTGCATCCACAAACGTGAGCTGACAGAACTGGACATCTACCACCGCATCCTCCGTTTCCAGAACTACATGGTGGCACTGGTTAACAAATCCCTCCTGCCTCTGCGCTTCCGCCTGCCTGGCCTCGGGGAAGCTGTCTTCTTCACCCGTGGTCTCAAGTACAACTTTGAGCTGATCCTCTTCTGGGGACCTGGCTCTCTGTTTCTCAATGAATGGAGCCTCAAGGCCGAGTACAAACGTGGGGGGCAACGGCTAGAGCTGGCCCAGCGCCTCAGCAACCGCATCCTGTGGATTGGCATCGCTAACTTCCTGCTGTGCCCCCTCATCCTCATATGGCAAATCCTCTATGCCTTCTTCAGCTATGCTGAGGTGCTGAAGCGGGAGCCGGGGGCCCTGGGAGCACGCTGCTGGTCACTCTATGGCCGCTGCTACCTCCGCCACTTCAACGAGCTGGAGCACGAGCTGCAGTCCCGCCTCAACCGTGGCTACAAGCCCGCCTCCAAGTACATGAATTGCTTCTTGTCACCTCTTTTGACACTGCTGGCCAAGAATGGAGCCTTCTTCGCTGGCTCCATCCTGGCTGTGCTTATTGCCCTCACCATTTATGACGAAGATGTGTTGGCTGTGGAACATGTGCTGACCACCGTCACACTCCTGGGGGTCACCGTGACCGTGTGCAGGTCCTTTATCCCGGACCAGCACATGGTGTTCTGCCCTGAGCAGCTGCTCCGCGTGATCCTCGCTCACATCCACTACATGCCTGACCACTGGCAGGGTAATGCCCACCGCTCGCAGACCCGGGACGAGTTTGCCCAGCTCTTCCAGTACAAGGCAGTGTTCATTTTGGAAGAGTTGCTGAGCCCCATTGTCACACCCCTCATCCTCATCTTCTGCCTGCGCCCACGGGCCCTGGAGATTATAGACTTCTTCCGAAACTTCACCGTGGAGGTCGTTGGTGTGGGAGATACCTGCTCCTTTGCTCAGATGGATGTTCGCCAGCATGGTCATCCCCAGTGGCTATCTGCTGGGCAGACAGAGGCCTCAGTGTACCAGCAAGCTGAGGATGGAAAGACAGAGTTGTCACTCATGCACTTTGCCATCACCAACCCTGGCTGGCAGCCACCACGTGAGAGCACAGCCTTCCTAGGCTTCCTCAAGGAGCAGGTTCAGCGGGATGGAGCAGCTGCTAGCCTCGCCCAAGGGGGTCTGCTCCCTGAAAATGCCCTCTTTACGTCTATCCAGTCCTTACAATCTGAGTCTGAGCCCCTGAGCCTTATCGCAAATGTGGTAGCTGGCTCATCCTGCCGGGGCCCTCCACTGCCCAGAGACCTGCAGGGCTCCAGGCACAGGGCTGAAGTCGCCTCTGCCCTGCGCTCCTTCTCCCCGCTGCAACCCGGGCAGGCGCCCACAGGCCGGGCTAACAGCACCATGACAGGCTCTGGGGTGGATGCCAGGACAGCCAGCTCCGGGAGCAGCGTGTGGGAAGGACAGCTGCAGAGCCTGGTGCTGTCAGAATATGCATCCACAGAGATGAGCCTGCATGCCCTCTATATGCACCAGCTCCACAAGCAGCAGGCCCAGGCTGAACCTGAGCGGCATGTATGGCACCGCCGGGAGAGTGATGAGAGTGGAGAAAGCGCCCCTGATGAAGGGGGAGAGGGCGCCCGGGCCCCCCAGTCTATCCCTCGCTCTGCTAGCTATCCCTGTGCAGCACCCCGGCCTGGAGCTCCTGAGACCACCGCCCTGCATGGGGGCTTCCAGAGGCGCTACGGTGGCATCACAGATCCTGGCACAGTGCCCAGGGTTCCCTCTCATTTCTCTCGGCTGCCTCTTGGAGGGTGGGCAGAAGATGGGCAGTCGGCATCAAGGCACCCTGAGCCCGTGCCCGAAGAGGGCTCGGAGGATGAGCTACCCCCTCAGGTGCACAAGGTATAA |
| --- | --- |
